# Ferroptosis-Protective Membrane Domains in Quiescence

**DOI:** 10.1101/2023.04.27.538084

**Authors:** Amalia H. Megarioti, Alexandros Athanasopoulos, Dimitrios Koulouris, Bianca M. Esch, Manousos Makridakis, Vasiliki Lygirou, Martina Samiotaki, Jerome Zoidakis, Vicky Sophianopoulou, Bruno André, Florian Fröhlich, Christos Gournas

## Abstract

Quiescence is a common cellular state, required for stem-cell maintenance and microorganismal survival under stress conditions or starvation. However, the mechanisms promoting quiescence maintenance remain poorly known. Plasma membrane components segregate into distinct microdomains, yet the role of this compartmentalization in quiescence remains unexplored. Here, we show that Flavodoxin-like proteins (FLPs), ubiquinone reductases of the yeast eisosome membrane compartment, protect quiescent cells from lipid peroxidation and ferroptosis. Eisosomes and FLPs expand specifically in respiratory-active quiescent cells, and mutants lacking either show accelerated aging, defective quiescence maintenance, and accumulate peroxidized phospholipids with monounsaturated or polyunsaturated fatty acids (PUFA). FLPs are essential for the extra-mitochondrial regeneration of the lipophilic antioxidant ubiquinol. FLPs, alongside the Gpx1/2/3 glutathione peroxidases, prevent iron-driven, PUFA-dependent ferroptotic cell death. Our work is the first description of ferroptosis-protective mechanisms in yeast and introduces plasma membrane compartmentalization as an important factor for the long-term survival of quiescent cells.

## Introduction

Quiescence is a reversible, non-proliferating state, during which cells display very low metabolic activity^1^. In complex eukaryotes, stem-cells, neurons, and oocytes are examples of quiescent cells. Stem-cell quiescence is critical for tissue maintenance, while exit from quiescence is essential for normal metazoan development, sexual reproduction, wound healing and cancer progression^1^. Quiescence is also critical in microorganisms, as >98% of microbes exist in this non-growing state^1^, allowing survival for long periods^1^ until more favorable conditions permit growth recovery. In the yeast *Saccharomyces cerevisiae*, cells differentiate into quiescent and senescent populations following glucose depletion^2^. Quiescent yeasts remain poorly studied, despite recent important efforts to identify subcellular quiescence-specific predictors, including the formation of actin-bodies, microtube bundles, telomere hyperclusters, proteasome storage granules, stress granules etc^3–5^. Yet, many of these subcellular changes only correlate with growth arrest or quiescence and are not essential for quiescence entry/maintenance^3^. An important exception is the reorganization of the mitochondrial network morphology typically observed at the onset of oxidative phosphorylation^6^. Respiration is essential for quiescence development by supporting successive waves of internal-pH-dependent subcellular re-organizations before transition of the cytoplasm to a glass-state^7^. The role of the plasma membrane in quiescence, however, remains unexplored.

The plasma membrane performs essential roles in selective nutrient exchange and environmental sensing and communication. The protein and lipid constituents of the plasma membrane are highly compartmentalized^8^. In yeast, several plasma membrane domains have been identified^9^. The most studied is the *Membrane Compartment occupied by* the arginine transporter *Can1* (MCC)^10^ (Figure 1A). MCCs correspond to static furrow-like invaginations of the plasma membrane^11^, associate with a subcortical protein scaffold termed eisosome^12^, have distinct lipidic composition^13–15^ and display structural and functional analogies to mammalian caveolae^16^. A main component of eisosomes is Pil1, a highly-abundant, self-assembling BAR (Bin, Amphiphysin, Rvs)-domain protein shaping the invagination^17^. Pil1 is the main organizer of eisosomes, whereas the role of its Lsp1 paralog remains unknown^12^. Nce102 is an additional organizer of eisosomes^10, 18^. In the absence of Pil1 or Nce102, MCC/eisosome plasma membrane patches disassemble, and eisosome-resident proteins either become homogeneous at the plasma membrane or re-localize to the cytoplasm^10, 12, 18^. MCC-resident membrane proteins include tetraspan members of the Nce102/MARVEL and Sur7 families, and several nutrient transporters^9, 19^ (Figure 1A). Eisosomes host regulators of sphingolipid biosynthesis and membrane stress, participating in the spatial separation of components of signalling cascades, including TORC2^18, 20–22^. Eisosomes also act as ubiquitylation-protective membrane domains for specific nutrient transporters ^9, 23–26^. During short-term nutrient starvation, eisosomes recruit stress granule components and conceal transporters from endocytosis, allowing efficient recovery upon nutrient re-addition^23, 27–29^. Yet, the role of eisosomes in quiescent cells and long-term starvation has not been investigated.

**Figure 1.**
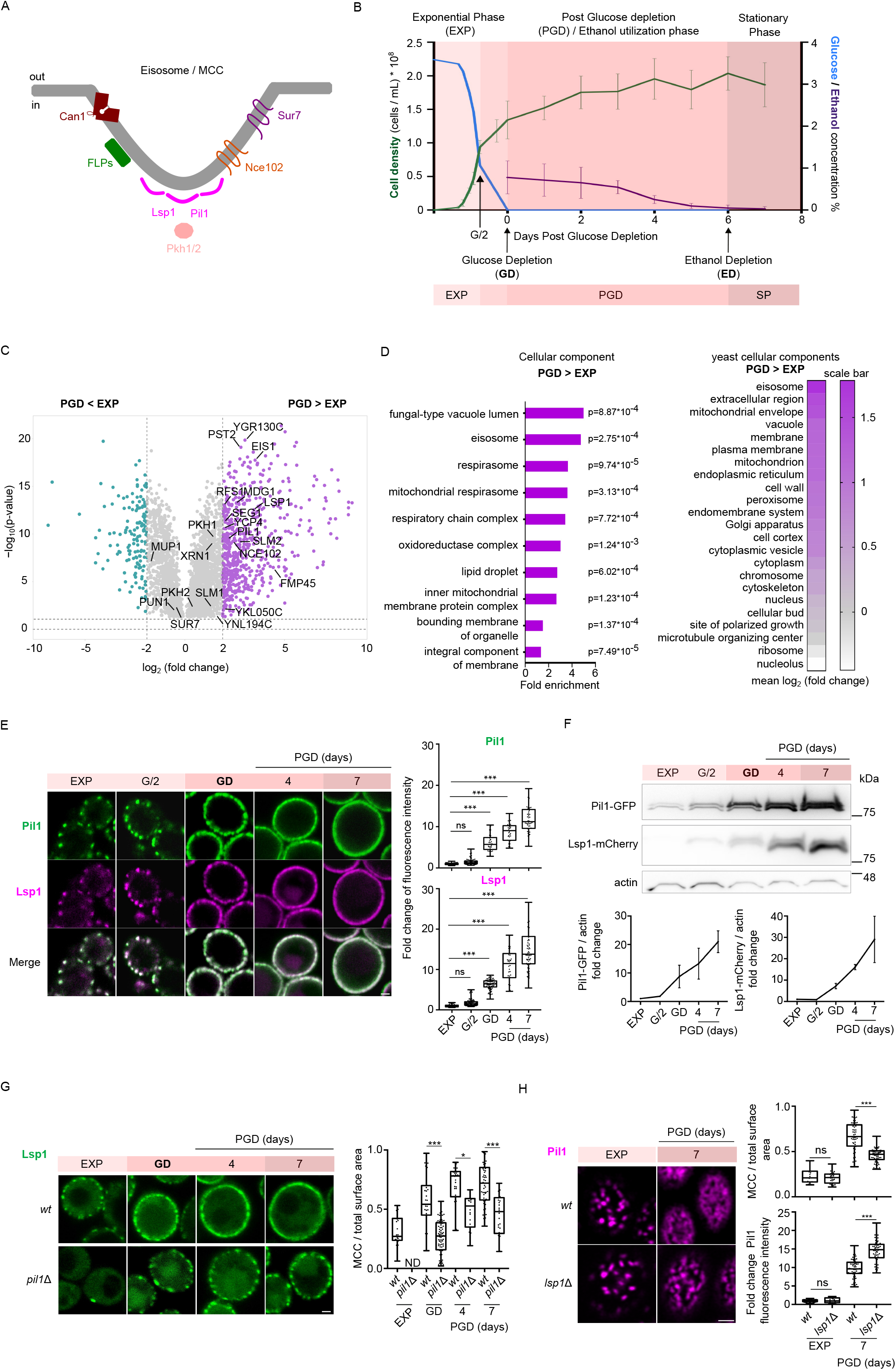
Eisosomes expand upon glucose depletion, and this requires the BAR-domain Lsp1 protein. **A)** Graphical representation of the *Membrane Compartment occupied by Can1* (MCC) which associates with a subcortical protein scaffold termed eisosome. The main components of eisosomes are annotated. **B)** Line-charts of cell density (green, cells/ml), glucose (blue) and ethanol (purple) concentrations (% w/v) at the different stages of cultures in minimal medium. Data from six (ethanol) or five (cell density) independent experiments are plotted over time as mean ± s.d. Glucose concentration of the average cell density per timepoint was calculated from the scatter-plot in Figure S1. Color-coding of the different growth phases is utilized throughout the manuscript. **C)** Volcano-plot [Student’s T-difference vs −log_10_(p-value)] comparison of the proteome of *wt* cells at GD vs EXP phase (Table S1). Proteins with significant > 1.9 (magenta) and < –1.9 (green) log_2_(fold-change) are indicated. All eisosomal proteins identified are annotated. **D)** Left: bar-chart of the top GO-TERM enriched cellular components in upregulated proteins of (C). The fold enrichment and p-values are shown. Right: heat map of the mean log_2_(fold-change) of all the proteins belonging to the indicated cellular components, and identified in (C). **E)** Middle section confocal microscopy images of a Pil1-GFP- and Lsp1-mCherry-expressing strain at different timepoints. Quantifications: Pil1-GFP and Lsp1-mCherry single-cell mean fluorescence intensities (n=35–97), normalized to the signal from EXP cells, are plotted in box and whiskers plots. \*\*\**P* < 0.001; ns, non-significant, p > 0.05, from one-way ANOVA. **F)** Western blot of total protein extracts from samples of (E), probed with anti-GFP, anti-mCherry and anti-actin. Quantifications: Signal intensities, normalized to actin, from 2 independent experiments are plotted in line-charts as mean fold-change ± s.d. from EXP samples. **G)** Middle section confocal microscopy of Lsp1-GFP-expressing *wt* and *pil1*Δ strains, at EXP and different timepoints PGD. Quantifications: The single-cell MCC/Total surface area ratios (n=20-84) from surface section images are plotted in box and whiskers plots. ND, not defined. \*\*\**P* < 0.001. \**P* < 0.05; ns, non-significant, p > 0.05, from one-way ANOVA. **H)** Surface section Airyscan confocal microscopy of Pil1-mCherry-expressing *wt* and *lsp1*Δ strains, at EXP and 7 days PGD. Quantifications: The single-cell MCC/Total surface area ratios and the Pil1-mCherry mean fluorescence intensities (n=18-59) normalized to the signal from EXP cells, from surface-section images are plotted in box and whiskers plots. \*\*\**P* < 0.001. ns, non-significant, p > 0.05, from one-way ANOVA. (Scale bar: 1 μm). See also Figure S1.

Ferroptosis is an iron-dependent form of non-apoptotic cell death in mammalian cells caused by increased peroxidation of plasma membrane polyunsaturated fatty acid (PUFA)-containing phospholipids^30–32^. The two main mechanisms protecting cells against ferroptosis are the reduction of lipid peroxides to lipid-alcohols by the GPX4 glutathione peroxidase^33^ and the inhibition of lipid-radical mediated autoxidation by ubiquinol, the reduced form of ubiquinone, generated by the Fsp1 reductase^34, 35^. In fungi, ferroptotic conidial cell death has been reported^36, 37^, but whether ferroptosis-protective mechanisms exist remains unknown. Yeast glutathione peroxidases Gpx1/2/3 are involved in protection from peroxidation^38, 39^, but whether this peroxidation is iron dependent has not been reported. Moreover, ubiquinone serves lipid peroxidation-protective roles in extra-mitochondrial membranes^40, 41^, but the ubiquinone reductases involved remain uncharacterized.

In this work, we identify an essential role of eisosomes in protecting quiescent yeasts from ferroptosis. Eisosomes expand in quiescent cells and stabilize the ubiquinone reductases, named FLPs^42^, which are involved in the extramitochondrial regeneration of ubiquinol in yeast. FLPs act in parallel with the glutathione peroxidases in protecting quiescent cells from detrimental lipid peroxidation and ferroptosis. The induction of both eisosomes and FLPs are requisites for the fitness and the long-term survival of quiescent yeasts.

## Results

### Eisosomes expand upon glucose depletion, and this requires the BAR-domain Lsp1 protein

Yeasts cultivated in buffered minimal medium typically show distinct phases of growth. Preadapted cells first grew exponentially for 30 h until they consumed half of the glucose (G/2) (Figure 1B). The next 12 h, they grew slower until glucose depletion (GD). This phase was followed by a 6-day post-glucose-depletion (PGD) slow growth, corresponding to utilization via oxidative phosphorylation of the ethanol produced during fermentation (Figure 1B). Since quiescent yeasts are formed following glucose depletion and during respiratory growth^6^, the conditions above seem appropriate to study the formation of quiescent cells in minimal media, as confirmed below. Moreover, the growth deceleration observed at G/2 (Figure 1B) likely reflects cell preparation for quiescence entry, previously suggested to begin one division before glucose depletion^3, 43^. To identify proteins involved in the transition to quiescence, we compared the proteome of cells from exponential phase and upon glucose-depletion, and focused on pronounced changes in expression. This analysis identified 592 upregulated and 148 downregulated proteins (Figure 1C; Table S1). GO-term analysis of the PGD downregulated dataset identified an enrichment of proteins involved in mRNA translation, rRNA maturation, and biosynthesis of amino acids and nucleobases (Figure S1B), consistent with these cells being slow-growing and in the process of becoming quiescent. The PGD upregulated dataset showed enrichment in constituents of aerobic respiration, carbohydrate metabolism, glutathione metabolism and cellular oxidant detoxification (Figure S1B), in line with the PGD shift from fermentation to oxidative phosphorylation^3, 6^. Accordingly, GO-term enrichment analysis for cellular components identified vacuoles, mitochondria and lipid droplets (Figure 1D), which have known roles in quiescence/respiratory growth^6, 7, 44–46^. Besides these organelles, proteins of the eisosome cellular component showed the second highest enrichment (Figure 1D). Moreover, eisosomal proteins showed the highest mean fold-increase in the whole PGD proteome, compared to all yeast cellular components (Figure 1C and 1D). Thus, we decided to characterize the role of eisosomes in quiescence.

We verified that the abundance of the core eisosomal proteins Pil1, Lsp1 and Nce102 gradually increased upon glucose exhaustion and until ethanol depletion (Figure 1E, F and S1C). In fermenting cells, their expression and the density of eisosomes were low. A ∼4-fold increase in expression was observed upon glucose depletion, reminiscent of cells shifted to acute carbon starvation^27, 28^. Yet, this increase was rather minor compared to the ∼20-fold gradual increase observed during ethanol utilization, suggesting that respiration could be involved in eisosome expansion (analyzed below).

Next, we characterized eisosome assembly following glucose depletion. Similar to fermentation^10, 12, 18^, Pil1 and Nce102 proved essential, as the proportion of Lsp1-GFP foci at the PM was drastically reduced in *pil1*Δ or *nce102*Δ cells (Figure 1G and S1D). Nce102 seemed to promote eisosome-assembly via regulating Pil1 phosphorylation (Figure S1E), as in EXP^18^. Intriguingly, eisosome-like Lsp1 spots were detected at the plasma membrane of *pil1*Δ cells in PGD timepoints (Figure 1G). These Lsp1 spots colocalized with two MCC-resident proteins, namely Ynl194c^10^ (Figure S1F), and the Can1(7KR) variant resistant to ubiquitylation and endocytosis^23^ (Figure S1G), suggesting that they correspond to intact eisosomes. Given that Lsp1 is neither necessary nor sufficient for eisosome formation during fermentation^12^ (Figure 1G), but can form furrows during stress^47^, these results indicate that Lsp1 exerts its main physiological role in quiescence. Consistently, we could observe an important reduction of the MCC/Total surface area in Lsp1-lacking quiescent cells, which was not due to reduced Pil1 levels (Figure 1H).

The above results show that eisosomes are among the most extensively expanding cellular components upon shift from fermentation to respiration, and identify Lsp1 as an additional, quiescence-specific eisosome organizer.

### Eisosomes expand in respiratory-active quiescent yeasts

Following glucose depletion, cells displayed heterogenicity regarding eisosome expansion and assembly. The majority showed eisosome expansion, yet a small percentage displayed eisosome disassembly and/or defective eisosome expansion (Figure 2A). We, thus, examined whether this heterogeneity correlates with the previously reported formation of quiescent and senescent cells in this phase^2^. Density-gradient enrichment of quiescent cells^2^ revealed that eisosome expansion occurred mainly in dense cells (Figure S2A and S2B). The less-dense, senescent-enriched fraction^2^ was heterogeneous and enriched in cells with defective eisosome expansion/assembly, but nevertheless contained cells with normal eisosome expansion (Figure S2A and S2B). These observations are consistent with recent reports suggesting that cell density cannot accurately discriminate quiescent and senescent cells^6^. Thus, we turned to the more refined marker of quiescence, namely the vesicular morphology of the mitochondrial network^6^ at single-cell level, and examined at different timepoints post-glucose depletion if it correlates with Pil1 induction and eisosome assembly (Figure 2A and 2B). Mitochondrial distribution changed from network-like in fermenting cells to vesicular (respiratory-competent) in 90% of cells at the beginning of ethanol utilization, indicative of quiescence^6^ (Figure 2B and 2C). The PGD population further contained 7% of apparently-senescent cells displaying globular mitochondria and 3% of non-fluorescent cells, reportedly dead^6^. Both non-quiescent categories increased overtime. Notably, quiescent cells were mostly homogeneous and the vast majority displayed normal eisosome expansion and peripheral Pil1 distribution (Figure 2B, 2C and S2C; Table S2). On the contrary, senescent cells were heterogeneous regarding eisosome expansion/assembly, with the majority, however, showing defective induction and cytoplasmic redistribution of Pil1, while intermediate situations were also detected (Figure 2B, 2C and S2C; Table S2). Finally, dead cells showed mostly eisosome disassembly. Inversely, eisosome expansion showed very high correlation with vesicular mitochondria (Figure 2D; Table S3). A cell with induced Pil1 had more than 96% probability of being quiescent until ethanol depletion. Moreover, cells showing both proper eisosome expansion and assembly had more than 89% probability of being quiescent at all timepoints. Although it was difficult to predict the fate of cells without Pil1 induction at early timepoints, these cells eventually tended to be mostly senescent or dead. These results indicate that the state of eisosomes can be a reliable marker of cell fate.

**Figure 2.**
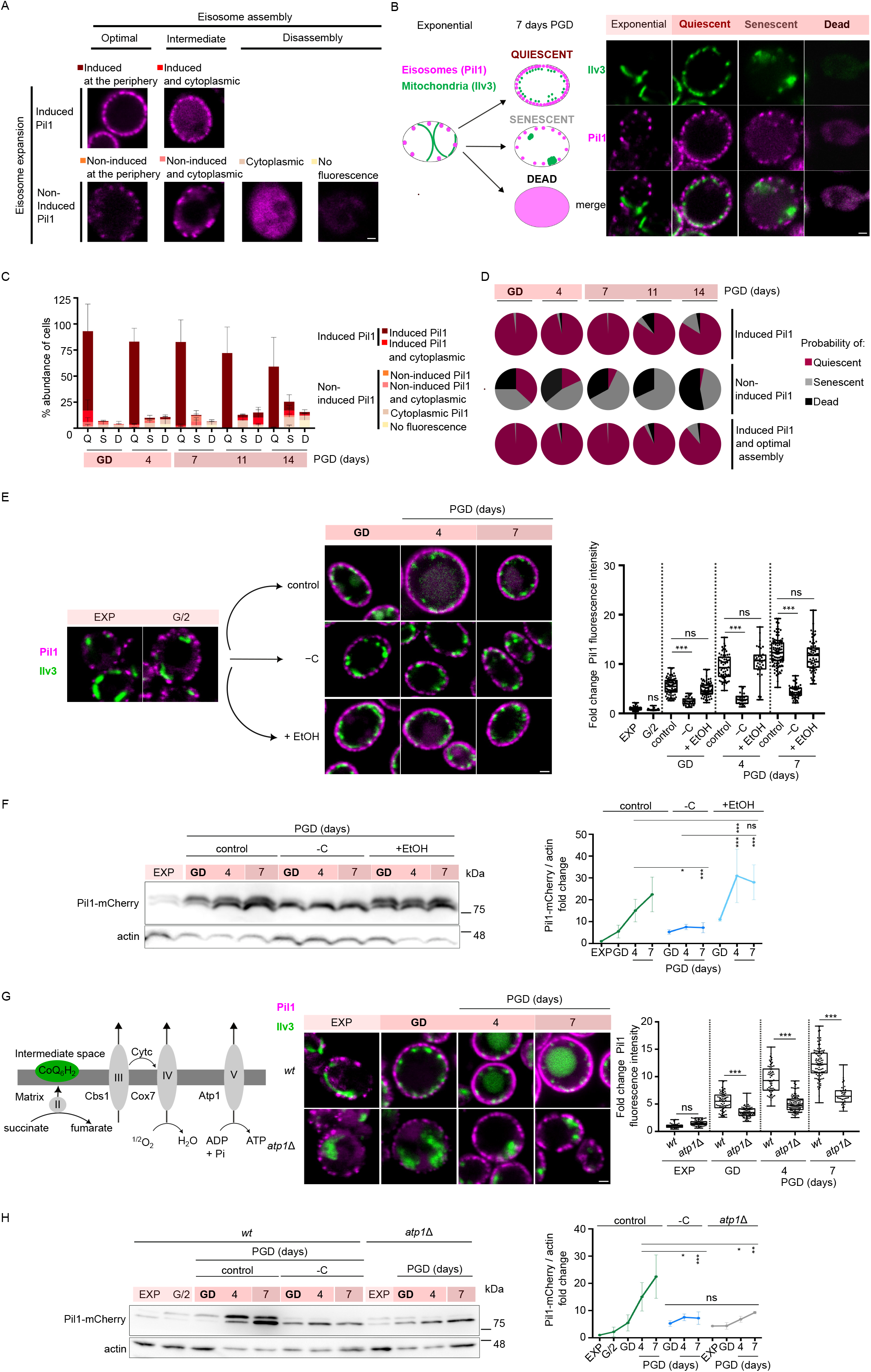
Eisosomes expand in respiratory-active quiescent yeasts. **A)** Middle section confocal microscopy images of a Pil1-mCherry-expressing strain, representative of all cell categories identified regarding eisosome assembly and expansion. **B)** Left: Schematic representation of the distribution of mitochondria according to^6^ and the correlation between mitochondrial distribution and eisosome-expansion/assembly at different timepoints PGD. Right: middle section confocal microscopy images of a Pil1-mCherry and Ilv3-GFP-expressing strain, representative of the predominant categories identified. **C)** Stacked bar-chart displaying the classification of cells from different timepoints PGD, according to mitochondrial distribution (as in B) and the eisosome-induction and/or assembly (as in A). Data from 4 independent experiments are represented as mean ± s.d. (detailed in Table S2, along with Chi-squared values), representative quantifications of Pil1 intensity are provided in Figure S2C. **D)** Pie charts illustrating the probability of a cell with optimal eisosome assembly and with or without Pil1 induction to be Quiescent, Senescent or Dead (according to B). Detailed calculations are provided in Table S3. **E)** Middle section confocal microscopy of a Pil1-mCherry- and Ilv3-GFP-expressing strain, at different timepoints PGD (control), or shifted at G/2 to glucose-free medium, supplemented with 1,18% w/v Ethanol (+EtOH) or not (–C). Quantifications: Pil1-mCherry single-cell fluorescence intensities (n=23-109), normalized to the signal from EXP cells, are plotted in box and whiskers plots. \*\*\**P* < 0.001; ns, non-significant, p > 0.05, from one-way ANOVA. **F)** Western blot of total protein extracts from a Pil1-mCherry-expressing strain, collected at the timepoints and conditions of (E), probed with anti-mCherry and anti-actin. Quantifications: Signal intensities, normalized to actin, from 2-4 independent experiments are plotted in line charts as mean fold-change ± s.d. from EXP samples. \*\*\**P* < 0.001; \*\**P* < 0.01; \**P* < 0.05; ns, non-significant, p > 0.05, from two-way ANOVA. **G)** Right: Middle section confocal microscopy of Pil1-mCherry- and Ilv3-GFP-expressing *wt* or *atp1*Δ strains, at EXP and different timepoints PGD, Quantifications: Pil1-mCherry single-cell fluorescence intensities (n=36-118), normalized to the signal from EXP *wt* cells, are plotted in box and whiskers plots. \*\*\**P*< 0.001; \*\**P* < 0.01; \**P* < 0.05; ns, non-significant, p > 0.05, from one-way ANOVA. Left: schematic representation of the mitochondrial respiratory chain. The respiratory complexes and the corresponding mutants are indicated. **H)** Western blot of total protein extracts from Pil1-mCherry-expressing *wt* or *atp1*Δ strains, collected at the timepoints and conditions of (G), probed with anti-mCherry and anti-actin. Quantifications: Signal intensities, normalized to actin, from 2-4 independent experiments are plotted in line charts as mean fold-change ± s.d. from *wt* EXP samples. \*\*\**P* < 0.001; \*\**P* < 0.01; \**P* < 0.05; ns, non-significant, p > 0.05, from two-way ANOVA. (Scale bar: 1 μm). See also Figure S2.

In support of eisosomes expanding specifically in respiratory-active cells, shifting cells from G/2 to fresh, carbon-free medium provoked only a minor induction in Pil1 expression (Figure 2E and 2F), as previously reported^27^. This indicates that glucose starvation is insufficient to promote optimal eisosome expansion. Consistently, the formation of stress-granules, visible upon acute glucose starvation^27^, did not increase PGD, while cells displaying stress-granules were mostly senescent, based on Pil1 expression/localization (Figure S2D). Antithetically, full eisosome expansion PGD required both glucose depletion and the supplementary presence of ethanol (Figure 2E and 2F), or another non-fermentable carbon source (Figure S2E). The requirement of respiration for eisosome expansion was further confirmed by the defective PGD Pil1 induction in mutants lacking essential components of respiratory chain complexes II, III and IV and *rho0* cells (Figure 2G, 2H, S2F and S2G).

In total, the above results strongly indicate that eisosomes are specifically induced in respiratory-active quiescent cells, while optimal eisosome expansion and assembly correlate strongly with quiescence maintenance.

### Eisosomes promote respiratory growth and are required for the long-term quiescence maintenance

Next, we focused on identifying the physiological role of eisosomes in quiescent cells, using the double *pil1*Δ*lsp1*Δ (*mcc*Δ) mutant which totally lacks eisosomes in PGD timepoints (Figure S1G). The *mcc*Δ strain grew identically to the *wt* during fermentation. However, this mutant displayed a more pronounced slowdown at G/2, and it showed limited growth thereafter (Figure 3A), a phenotype intermediate between the *wt* and the *atp1*Δ respiration-deficient strain. This defective PGD growth could be largely complemented by ectopic expression of either *PIL1* or *LSP1* (Figure S3A), confirming that the phenotype is due to the absence of eisosomes. The above suggest that *mcc*Δ cells might suffer from defective PGD respiratory metabolism. Indeed, cultures of the *mcc*Δ strain showed reduced consumption of ethanol, similarly to a respiration-deficient mutant (Figure 3B). This impaired respiratory growth was not due to defective induction of autophagy (Figure S3B), which is essential for adaptation to respiration^48^. Rather, *mcc*Δ cells displayed defective mitochondrial functionality following glucose depletion, and more specifically problematic mitochondrial membrane potential (MMP). More specifically, *mcc*Δ cells showed defective targeting of PreCOX4-mCherry to mitochondria, a process known to require intact MMP^49^, comparably to *wt* cells treated with the MMP-uncoupler CCCP (Figure 3C). This phenotype was also observed in cells with vesicular mitochondrial morphology. This indicates that, in the absence of eisosomes, mitochondria are not fully functional post-glucose depletion even in cells displaying a “respiratory-competent” phenotype.

**Figure 3.**
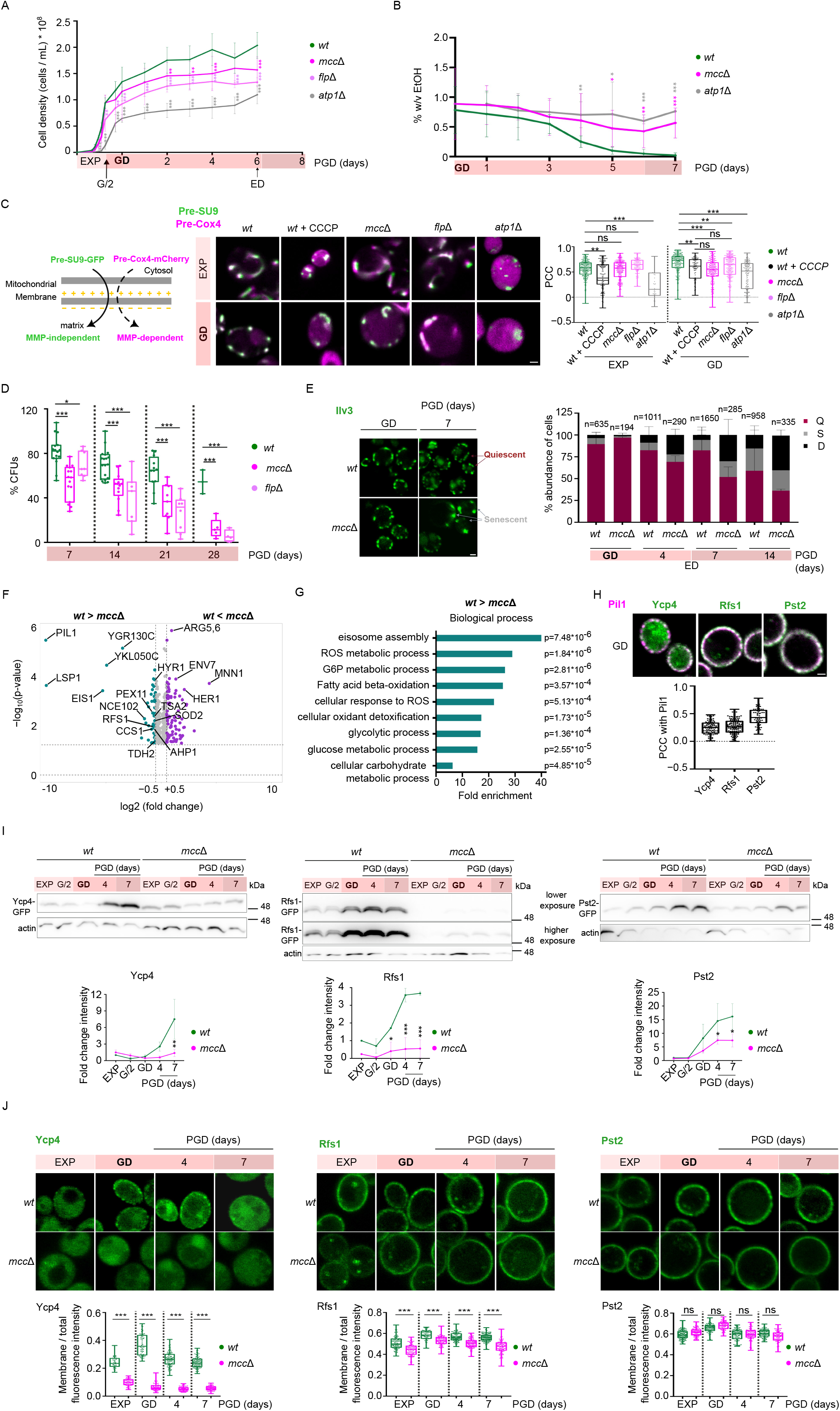
Eisosomes promote respiratory growth and are required for the long-term quiescence maintenance by stabilizing the Flavodoxin-Like Proteins. **A)** Line chart of *wt*, *pil1*Δ*lsp1*Δ (*mcc*Δ), *atp1*Δ and *pst2*Δ*ycp4*Δ*rfs1*Δ (*flp*Δ) cultures. Cell density measurements at different timepoints in EXP and PGD, from 5 independent experiments, are plotted as mean ± s.d. \*\*\**P* < 0.001; \*\**P* < 0.01; \**P* < 0.05, p > 0.05, from two-way ANOVA. **B)** % (w/v) ethanol measurements in supernatants of *wt*, *pil1*Δ*lsp1*Δ (*mcc*Δ) and *atp1*Δ cultures, from 4-6 independent experiments, at different timepoints PGD are plotted as mean ± s.d. \*\*\**P* < 0.001; \*\**P* < 0.01; \**P* < 0.05, from two-way ANOVA. **C)** Left: Schematic representation of the mitochondrial membrane potential (MMP) microscopy-based assay. *wt*, *pil1*Δ*lsp1*Δ (*mcc*Δ), *pst2*Δ*ycp4*Δ*rfs1*Δ (*flp*Δ) and *atp1*Δ strains are transformed with pMitoLoc plasmid^49^, encoding for Pre-SU9-GFP and Pre-Cox4-mCherry. Pre-SU9-GFP is targeted constitutively to the mitochondrial lumen, while targeting of Pre-Cox4-mCherry requires intact MMP^49^. Middle: Middle section merged confocal microscopy images of the indicated strains, treated or not for 6 hours with 30 μΜ CCCP and mutant strains, at EXP or at GD. Right: Quantifications: The single-cell Pearson correlation coefficient (PCC) of the two fluorescence signals (n=38-232), are plotted in box and whiskers plots. \*\*\**P* < 0.001; \*\**P* < 0.01; \**P* < 0.05; ns, non-significant, p > 0.05, from one-way ANOVA. **D)** Growth recovery of *wt*, *pil1*Δ*lsp1*Δ (*mcc*Δ) and *pst2*Δ*ycp4*Δ*rfs1*Δ (*flp*Δ) strains from different timepoints PGD, as percentages of CFUs from 3-19 independent experiments, are plotted in box and whiskers plots. \*\*\**P* < 0.001; \*\**P* < 0.01; \**P* < 0.05 from unpaired T-tests. **E)** Eisosomes are required for maintenance of quiescence. Middle section confocal microscopy images of Ilv3-GFP-expressing *wt* and *pil1*Δ*lsp1*Δ (*mcc*Δ) strains at GD and 7 days PGD (detailed in Figure S3D). Quantifications: the percentages *wt* and *pil1*Δ*lsp1*Δ (*mcc*Δ) cells displaying cortical (Quiescent, Q) or globular (Senescent, S) mitochondria, or lack of Ilv3-GFP fluorescence (Dead, D), (as in Figure 2B and 2C) at different timepoints PGD are plotted in stacked bar charts. Data from 2-4 independent experiments are represented as mean ± s.d. **F)** Volcano-plot [Student’s T-difference vs −log_10_(p-value)] comparison of the proteome of *pil1*Δ*lsp1*Δ (*mcc*Δ) vs *wt* cells at 3 days PGD (Table S4). Proteins with significant > 0.50 (magenta) and < –0.50 (green) log_2_(fold-change) are indicated. Eisosomal proteins, proteins related to ROS, and proteins differing most are highlighted. **G)** Bar graph of the top GO-TERM enriched biological processes in downregulated proteins of (F). The fold enrichment and p-values are shown. **H)** Middle section confocal microscopy merged images of strains expressing Pil1-mCherry and GFP-tagged versions of Ycp4, Rfs1 or Pst2, at GD. Quantifications: The single-cell Pearson correlation coefficient (PCC) values from surface sections of the two fluorescence signals (n=67-98), are plotted in box and whiskers plots. **I)** Western blot of total protein extracts from *wt* and *mcc*Δ strains expressing GFP-tagged versions of Ycp4, Rfs1 or Pst2, at EXP and different timepoints PGD probed with anti-GFP and anti-actin. Quantifications: Signal intensities, normalized to actin, from 2-3 independent experiments are plotted in line charts as mean fold-change ± s.d. from *wt* EXP samples. \*\*\**P* < 0.001; \*\**P* < 0.01; \**P* < 0.05, from two-way ANOVA. **J)** Middle section confocal microscopy from the strains and timepoints of (I). Quantifications: The single-cell membrane-to-total fluorescence intensity (n=40-117) are plotted in box and whiskers plots. \*\*\**P* < 0.001; ns, non-significant, p > 0.05, from one-way ANOVA. (Scale bar: 1 μm). See also Figure S3.

As mutants with defective respiration are known to age and die much faster^6^, we next sought to characterize the role of eisosomes in quiescent cell survival. In agreement with previous reports^2, 6^, growth recovery of the *wt* was close to 80% the first week PGD, and remained high for at least one month. On the contrary, the *mcc*Δ and other strains lacking MCC organizers, displayed only 50 % survival after the first week, and this dropped to 15 % within one month (Figure 3D and S3C). The reduced long-term survival was further confirmed at the single-cell level using the morphology of the mitochondrial network (Figure 3E and S3D). *mcc*Δ cells seemed fully capable of forming vesicular mitochondria following glucose depletion, indicating no defects in initial quiescent cell formation. However, the percentage of quiescent cells containing vesicular mitochondria declined overtime much faster than the *wt*. The mutant population acquired significantly increased percentages of both senescent cells containing globular-mitochondria, and non-fluorescent dead cells (Figure 3E and S3D). Overall, these results show that eisosomes support respiratory growth after glucose depletion and are required for the long-term quiescence maintenance and growth recovery.

### Eisosomes promote quiescence maintenance by stabilizing the Flavodoxin-Like Proteins

For characterizing how eisosomes affect quiescence maintenance, we compared the proteome of *wt* and *mcc*Δ cells (Figure 3F; Table S4) during the ethanol utilization phase (Figure 1B). This analysis identified 59 downregulated and 90 upregulated proteins in the mutant (Table S4). 58% of the downregulated proteins overlap with the PGD-upregulated proteins (Figure S3E), suggesting partially defective adaptation to quiescence. Indeed, proteins involved in important quiescence-related biological processes^3^ (Figure 3G), such as respiration-related metabolism of carbohydrates and fatty acids, were enriched in the downregulated proteins of the *mcc*Δ strain. As expected, the top category identified was eisosome assembly. Intriguingly, the dataset of proteins downregulated in the *mcc*Δ was enriched in proteins involved in Reactive Oxygen Species (ROS) metabolism and detoxification. The downregulation of these processes after the post-glucose depletion shift to respiration, when increased mitochondrial ROS production occurs^7^, could contribute to the defective quiescence preservation of the *mcc*Δ (Figure 3E). Our attention was drawn by Rfs1, an eisosome-resident peripheral membrane protein with potential antioxidant role^10, 42^. Rfs1 is one of the three (along with Pst2 and Ycp4) poorly characterized Flavodoxin-Like Proteins (FLPs) of yeast^10^. All three FLPs were upregulated PGD (Figure 1C), suggesting an important quiescence-specific role. As during fermentation^10^, FLPs showed mainly eisosomal localization post-glucose depletion (Figure 3H), while they were many-fold induced in quiescent cells (Figure 3I). Ycp4 displayed cytoplasmic and eisosomal localization, while Pst2 and Rfs1 were found at the plasma membrane and showed enrichment in eisosomes (Figure 3H and 3I). Most importantly, in *mcc*Δ cells, all three FLPs were significantly less abundant (Figure 3I), Pst2 and Rfs1 became homogeneous at the plasma membrane, while Rfs1 and especially Ycp4 additionally showed defective recruitment to the plasma membrane (Figure 3J). Although the mechanism of plasma membrane/eisosomal recruitment of FLPs is unknown, Ycp4 is palmitoylated^50^ and this modification could assist its tethering to membranes. The above results show that the total levels of FLPs at the plasma membrane of the *mcc*Δ strain are much reduced. To determine whether the phenotypes of the *mcc*Δ strain could be attributed to the lower levels of FLPs, we examined the phenotype of the FLP-lacking (*flp*Δ) strain. The *flp*Δ not only phenocopied the *mcc*Δ for defective PGD growth, long-term survival and mitochondrial membrane potential, but also showed stronger defects (Figure 3A, 3C and 3D). These defects were not due to impaired eisosome assembly (Figure S3F). Consistently, the PGD proteome of the *flp*Δ strain showed, similarly to the one of the *mcc*Δ, defective induction of proteins involved in mitochondrial respiration and the metabolism of carbohydrates and fatty acid acids (Figure S3G and S3H; Table S4). We conclude that stabilization of the FLPs at the plasma membrane is an important function of eisosomes for promoting long-term quiescence maintenance.

### Eisosomes and Flavodoxin-Like Proteins protect quiescent cells from lipid peroxidation and ferroptosis

Next, we characterized how the lack of FLPs affects quiescent cells. Pst2 shows reductase activity against several quinones, including ubiquinone^51^. The reduced form of ubiquinone, ubiquinol, is part of the respiratory chain but also a lipophilic antioxidant protecting yeasts from lipid peroxidation, with roles in extra-mitochondrial membranes^40, 41^. We hypothesized that FLPs could act analogously to the plasma membrane-localized ubiquinol-generating Fsp1 reductase, which protects human cells from ferroptosis^34, 35^. Consistently, *mcc*Δ cells and especially *flp*Δ cells were more sensitive than the *wt* to supplementation with polyunsaturated fatty acids (PUFAs) (linolenic acid, LNA; linoleic acid, LA), which are more prone to peroxidation^52^, in respiratory media (Figure 4A and S4A). Among the FLPs, deletion of *PST2* caused the most pronounced effect, while additional deletion of *RFS1* and *YCP4* further enhanced LNA sensitivity (Figure S4B). The sensitivity of the *mcc*Δ to LNA could be complemented by ectopic expression of *PIL1,* and to a lesser extent *LSP1* (Figure S4C). Hypersensitivity was more evident to PUFAs than monounsaturated (MUFAs) or saturated fatty acids (Figure S4A), pointing to accumulation of peroxidized phospholipids at the plasma membrane of the mutants. To directly measure peroxidized phospholipids, we performed targeted lipidomics of PGD cells, from cultures supplemented or not with either MUFAs (oleic acid) or PUFAs (LA or LNA). We determined the peroxidation of the plasma membrane-enriched phospholipid Phosphatidyl-Ethanolamine (PE) as a readout, since PE can incorporate exogenous fatty acids to its backbone^53^, while peroxidation of PE is considered the main driver of ferroptosis in human cells^54^. Indeed, externally-supplied MUFAs and PUFAs were incorporated in the PE of all strains to similar levels (Figure 4B). Most importantly, two days PGD, we could identify a several-fold increase in peroxidized PUFA-containing PE, specifically in *mcc*Δ and *flp*Δ cells grown in the presence of PUFAs (Figure 4C and 4D). These results strongly indicate an important role of FLPs and ubiquinol in counteracting lipid peroxidation, mainly in respiratory conditions and quiescent cells.

**Figure 4.**
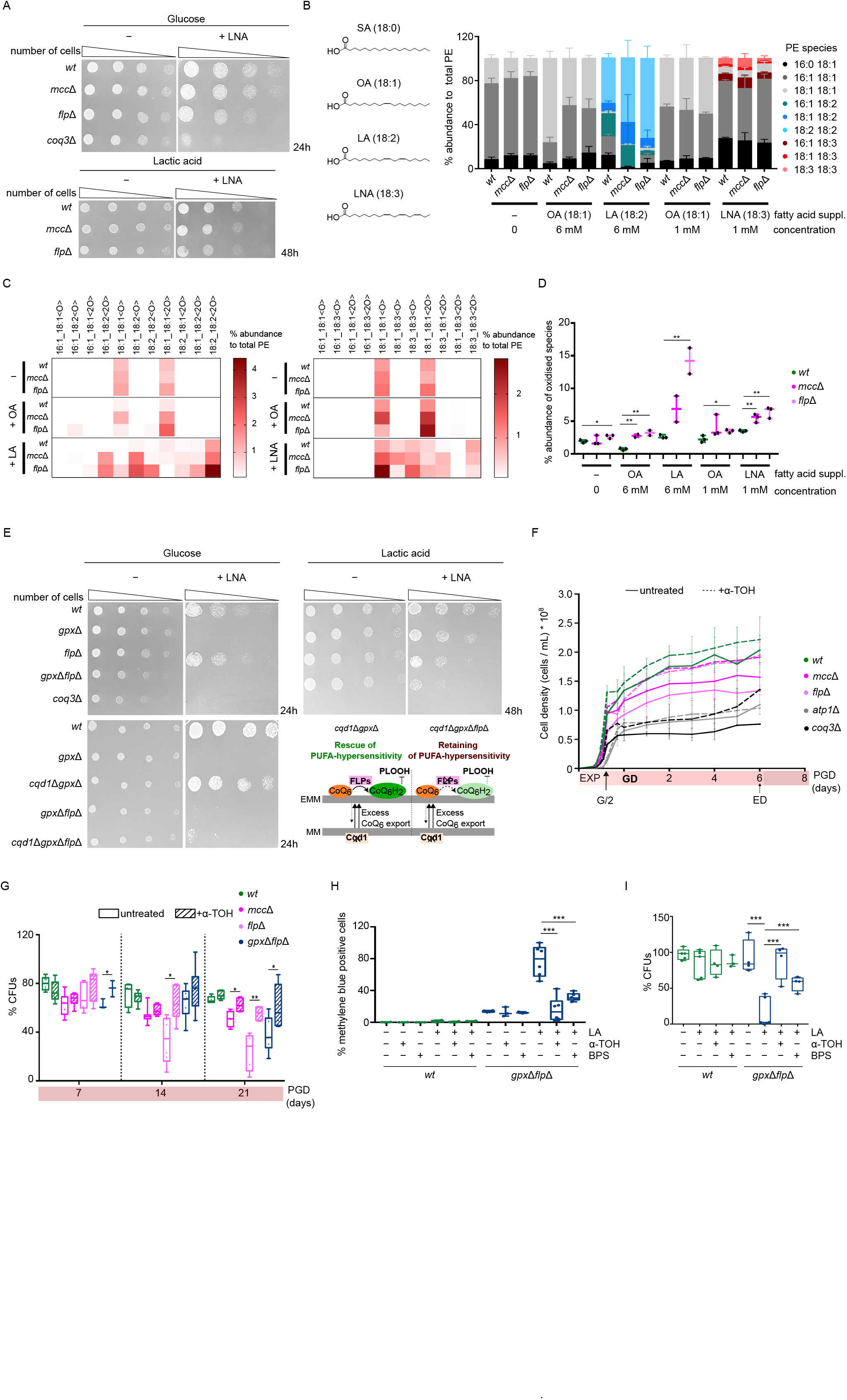
Eisosomes and FLPs protect Quiescent cells from lipid peroxidation and ferroptosis. **A)** Drop-tests of *wt*, *pil1*Δ*lsp1*Δ (*mcc*Δ), *pst2*Δ*ycp4*Δ*rfs1*Δ (*flp*Δ) and *coq3*Δ strains in minimal medium with glucose or lactic acid as carbon source, supplemented or not with 0.1 mM linolenic acid (LNA). **B)** Incorporation of exogenously-supplied fatty acids, at indicated concentrations, in phosphatidylethanolamine (PE) of *wt, mcc*Δ, and *flp*Δ cells at 2 days PGD. Each PE species was identified using multiple reaction monitoring as shown in Table S6. The % abundance of each PE species normalized to the total PE detected in each sample from 2-3 biological replicates and 2 technical replicates are plotted in stacked bar charts as mean ± s.d. **C)** Heat map of the % abundance of each oxidized (<O>) and peroxidized (<2O>) PE species normalized to total PE in PUFA (LA, LNA) and MUFA (OA) supplemented and unsupplemented *wt*, *mcc*Δ and *flp*Δ cells. The coloring threshold was set to 0.1% abundance. **D)** The % abundance of the total oxidized and peroxidized PE detected in PUFA (LA, LNA) and MUFA (OA) supplemented and unsupplemented *wt*, *mcc*Δ and *flp*Δ cells are plotted as box and whiskers. \*\*\**P* < 0.001; \*\**P* < 0.01; \**P* < 0.05; ns, non-significant, p > 0.05, from unpaired T-tests. **E)** Drop-tests of *wt*, *gpx1*Δ*gpx2*Δ*gpx3*Δ (*gpx*Δ), *pst2*Δ*ycp4*Δ*rfs1*Δ (*flp*Δ), *gpx*Δ*flp*Δ, *coq3*Δ, *cqd1*Δ*gpx*Δ, and *cqd1*Δ*gpx*Δ*flp*Δ strains in minimal medium with glucose or lactic acid as carbon source, supplemented or not with 0.1 mM linolenic acid (LNA). Scheme: Graphical representation of the interpretation of the drop-tests. Cqd1 is involved in import of ubiquinone (CoQ6) in mitochondria and *cqd1*Δ rescues the PUFA-hypersensitivity of *gpx*Δ cells, by excess ubiquinone export from the mitochondrial membrane (MM) to extra-mitochondrial membranes (EMM)^40^. Additional lack of FLPs is epistatic over Cqd1-dependent rescue of PUFA-hypersensitivity of *gpx*Δ cells, indicating that FLPs are required for extra-mitochondrial ubiquinol (CoQ_6_H_2_)-mediated protection from lipid peroxidation. **F)** Line-charts of *wt*, *pil1*Δ*lsp1*Δ (*mcc*Δ), *pst2*Δ*ycp4*Δ*rfs1*Δ (*flp*Δ), *atp1*Δ and *coq3*Δ yeast cultures in minimal medium, split and supplemented or not with 0.5 mM α-tocopherol (+α-TOH) at OD_600_=3. Cell density measurements at different timepoints in EXP and PGD, from 5 independent experiments are plotted as mean ± s.d. Statistical analysis from two-way ANOVA is provided in Table S5. **G)** Growth recovery of *wt*, *pil1*Δ*lsp1*Δ (*mcc*Δ), *pst2*Δ*ycp4*Δ*rfs1*Δ (*flp*Δ) and *gpx*Δ*flp*Δ yeast cultures in minimal medium supplemented or not with 1 mM α-tocopherol (+α-TOH) at OD_600_=3, as percentages of CFUs from 3-7 independent experiments, are plotted in in box and whiskers plots. \*\*\**P* < 0.001; \*\**P* < 0.01; \**P* < 0.05, from unpaired T-tests. **H and I)** % of dead, methylene blue-stained cells (H) or growth recovery (I) at 2 days PGD of *wt* and *gpx*Δ*flp*Δ strains, treated or not with 10 mM Linoleic acid (LA) and 1 mM α-tocopherol (α-TOH) or 100 μΜ iron chelator (BPS) at OD_600_=3. Data from 3-6 independent experiments are plotted in box and whiskers plots. \*\*\**P* < 0.001; \*\**P* < 0.01; \**P* < 0.05. ns, non-significant, p > 0.05, from unpaired T-tests. See also Figure S4.

Consistent with the above view, *flp*Δ and *mcc*Δ cells showed only limited sensitivity to LNA in glucose media and were less sensitive to LNA than the *coq3*Δ strain (Figure 4A), which totally lacks ubiquinone and is hypersensitive to lipid peroxidation^41^. This mild phenotype could be explained by a contribution of the Gpx1/2/3 glutathione peroxidases reported to counteract lipid peroxidation during fermentation^39^. Indeed, a strain carrying deletions of the *GPX1/2/3* (*gpx*Δ) showed hypersensitivity to LNA in glucose but was less sensitive to LNA than the *flp*Δ in respiratory conditions (Figure 4E). These results suggest that Gpx1/2/3-glutathione is the principal lipid-ROS scavenging mechanism during fermentation, while during quiescence FLPs-ubiquinol have a more prominent role. In support of FLPs acting in parallel with the Gpx1/2/3 to protect quiescent cells from peroxidation, a strain carrying deletion of all six genes (*gpx*Δ*flp*Δ) displayed synthetic hypersensitivity in respiratory conditions (Figure 4E). We further confirmed genetically that FLPs were essential for the extra-mitochondrial role of ubiquinone in counteracting lipid peroxidation, by deleting *CQD1,* which is required for mitochondrial import of ubiquinone^40^ (Figure 4E). Lack of Cqd1 results in excess ubiquinone export to extra-mitochondrial membranes and rescues the hypersensitivity of the *gpx*Δ strain to LNA^40^. Importantly, FLPs were epistatic over this Cqd1-dependent rescue of the LNA hypersensitivity of the *gpx*Δ (Figure 4E). Thus, this result identifies FLPs as the main extra-mitochondrial ubiquinol regenerating reductases.

The results above highlighted the importance of FLPs in counteracting lipid peroxidation in PUFA-supplemented cultures, consistent with PUFAs being more prone to peroxidation^52^. Although yeast does not biosynthesize PUFAs, our results (Figure 3) reveal an essential role of eisosomes and FLPs in PUFA-unsupplemented quiescent cells, suggesting that the phenotypes could be caused by peroxidation of MUFAs. Indeed, we could detect a small but significant increase in peroxidized PE-incorporated MUFA phospholipids for the *flp*Δ mutant in unsupplemented cultures (Figure 4C and 4D). Exogenous supplementation with MUFAs led to significantly increased levels in PE-MUFA peroxides in both the *flp*Δ and *mcc*Δ cells. In further support of MUFA-containing phospholipid peroxidation as the main defect of the eisosome- and FLP-lacking strains, supplementation of the cultures with the lipid peroxidation-scavenger α-tocopherol^41^ could significantly rescue the defects in PGD growth of these mutants (Figure 4F; Table S5). Significant α-tocopherol-rescue was also observed in ubiquinone-lacking *coq3*Δ mutants but not in *wt* or *atp1*Δ respiration-deficient cells, indicating that the rescue concerns peroxidation-protective roles of ubiquinone and not respiratory- chain-related roles. Importantly, supplementation with α-tocopherol was also sufficient to rescue the defective long-term growth recovery of the *mcc*Δ, *flp*Δ and *gpx*Δ*flp*Δ strains (Figure 4G).

The above findings strongly suggest that a mild accumulation of peroxidized MUFAs (Figure 4C and 4D) is responsible for the PGD growth-arrest, and defective quiescence maintenance observed in the mutants (Figure 3E, 4F and 4G). Yet, this accumulation seemed insufficient for causing rapid cell death and only led to reduced long-term survival. However, we could identify nearly 100% cell death, specifically for *gpx*Δ*flp*Δ cells, within two days of supplementation with 10 mM of the PUFA LA (Figure 4H and 4I). This PUFA-induced cell death was totally reversed by co-treatment with α-tocopherol, strongly pointing to peroxidation of lipids as the cause. Co-treatment with the Fe^2+^ chelator BPS could similarly largely rescue the PUFA-induced cell death, and to a lesser extent the growth recovery of the *gpx*Δ*flp*Δ (Figure 4H and 4I). This suggests that iron is essential for peroxidation of PUFA-containing phospholipids in quiescent yeasts.

We conclude that the eisosome-resident, ubiquinol-generating FLPs act in parallel with the Gpx1/2/3 for preventing quiescent yeast cell-death induced by peroxidation of PUFA- and MUFA-phospholipids (Figure 5). In the presence of exogenously-supplied PUFAs, lipid peroxidation seems iron-dependent, more pronounced and can lead to cell-death of a strain lacking the FLPs and the Gpx1/2/3, pointing out to the existence of ferroptosis in PUFA-supplemented quiescent yeasts.

**Figure 5.**
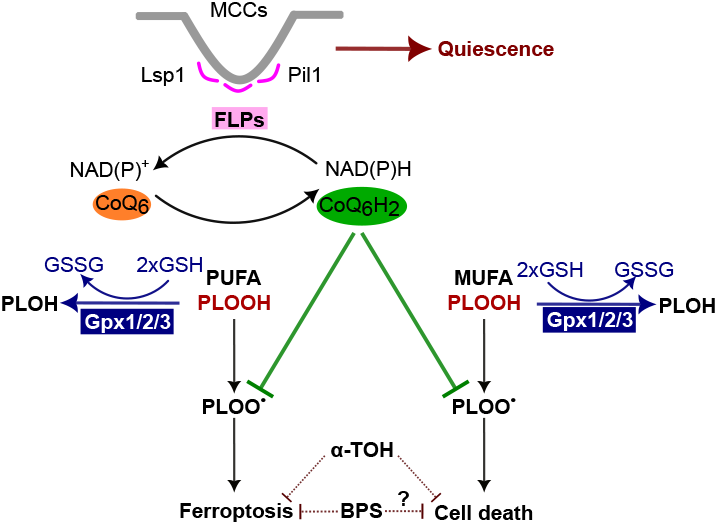
Eisosomes and FLPs protect quiescent yeasts from ferroptosis in parallel with the glutathione peroxidases. Schematic summary of the results. Eisosomes promote quiescence maintenance by stabilizing the Flavodoxin-like proteins (FLPs). FLPs counteract ferroptosis and cell death by regenerating ubiquinol (CoQ_6_H_2_), thereby inhibiting the lipid-radical (PLOO.) autooxidation of MUFAs and PUFAs^34, 35^, similarly to α-tocopherol (α-TOH). In parallel, the Gpx1/2/3 glutathione (GSH) peroxidases reduce peroxidized MUFAs and PUFAs (PLOOH) to lipid alcohols (PLOH)^33, 39^. The iron chelator (BPS) rescues from ferroptosis in PUFA-supplemented cultures.

## Discussion

In this work, we identify MCCs/eisosomes as a highly-expanding plasma membrane compartment in quiescence. We further show that eisosomes are essential for quiescence maintenance and the long-term survival of quiescent cells. The ubiquinol-generating^51^, eisosome-resident FLPs, in particular, protect quiescent yeasts from phospholipid peroxidation. The FLP Pst2 affects the survival in stationary phase^55^, but the mechanism remained unknown. Our results are consistent with the high abundance of eisosomes in quiescent conidiospores and ascospores of *Aspergillus nidulans*^56, 57^, the peroxidation-protective roles and the increased levels of ubiquinol in stationary phase^40, 41, 58^, and the known antioxidant roles of FLPs in *Candida albicans* against exogenous quinones and PUFAs^42, 59^. The absence of eisosomes causes similar but less severe phenotypes than the lack of FLPs (Figure 3A, 3C and 3D), consistent with the reduced membrane abundance/recruitment of FLPs in the *mcc*Δ (Figure 3J). Yet, it cannot be excluded that eisosomes have additional, FLP-independent roles in protection from lipid peroxidation. For example, the abundance of Gpx3/Hyr1, which reduces hydroperoxy-phospholipids^39^ and contributes to protection from ferroptosis (Figure 4), is reduced in quiescent *mcc*Δ cells (Figure 3F), but the underlying mechanism remains elusive. Other properties of eisosomes, like membrane curvature, the particular lipidic composition^13–15^ and TORC2-signalling^20, 21^, could additionally affect protection from peroxidation. Interestingly, our proteomics data suggest previously unnoticed interactions between eisosomes and fatty-acid oxidation (Figure 3G and S3H), a process potentially involved in sensitivity to lipid peroxidation^60^.

In the presence of PUFAs, we detected increased levels of peroxidized PUFA-containing PE, which is considered a main driver of ferroptosis in human cells^54^. PUFA-mediated phospholipid peroxidation seems iron-dependent and can lead to cell death of a strain lacking both the Gpx1/2/3 and the FLPs (Figure 4H and 4I). These two mechanisms, namely the one of glutathione and glutathione peroxidases and the one of ubiquinol and FLPs, act in parallel to protect yeasts from peroxidation under different physiological conditions (Figure 4E). The analogy with Gpx4 and Fsp1 acting in parallel in human cells^34, 35^ strongly indicates that ferroptosis-like mechanisms are conserved in fungi, and they are important for quiescent fungal cell survival. As quiescent yeasts are critical for environmental spreading^61^, ferroptosis-protective mechanisms could be essential in linoleic acid containing natural yeast habitats, like grapes. Ferroptosis-protective mechanisms could additionally have important roles in other, PUFA-biosynthesizing/incorporating fungi, including pathogenic fungi. Interestingly, we have identified homologues of human Fsp1 in several PUFA-biosynthesizing filamentous fungi, including 2 homologues in *A. nidulans*, which are under investigation (VS, CG, unpublished results). Quiescent yeasts are implicated in the development of persistence, a resistance to antimicrobial compounds targeting only growing cells^62^. Given that eisosomes are fungal-specific structures and abundantly expressed in dormant cells/spores of other fungal species as well^9, 56, 57, 63^, eisosomal proteins could be promising targets for the development of novel, growth-independent fungal treatment strategies.

An important aspect of our work is the link of MUFA peroxidation with survival defects. MUFA-peroxidation is considered less efficient than PUFA-peroxidation, while MUFA supplementation promotes a ferroptosis-resistant state^52, 60^. Surprisingly, our results indicate that MUFA-peroxidation can occur in the absence of PUFAs and can lead to growth defects that are rescued by addition of α-tocopherol (Figure 4C, 4D, 4F and 4G). Yet, MUFA-peroxidation seems less efficient than PUFA-peroxidation and does not cause rapid cell death (Figure 4C, 4D, 4F and 4G). Determining whether MUFA peroxidation is iron-dependent is not feasible under current experimental conditions, since iron chelation would affect respiration and consequently quiescence. Interestingly, other transition metals have previously been connected to lipid peroxidation in yeast^38^. Fungal FLPs are distantly related to the WrbA bacterial and NQO1/2 mammalian proteins, which have known ubiquinol-generating activities^64^. Yet, the involvement of NQO1/2 in protection from ferroptosis has only occasionally been mentioned^65, 66^, while less is known about the bacterial proteins^30^. Future work could address whether NQO1/2 have condition/cell/organelle-specific roles in counteracting ferroptosis, or whether WrbA proteins have lipid peroxidation-protective roles in PUFA-containing/incorporating bacteria.

Lipid peroxidation in human cells has been previously connected to defective mitochondrial function, but the causal relationship remained controversial and context-dependent^67, 68^. In our conditions, α-tocopherol supplementation rescued the defects in PGD growth and growth recovery (Figure 4F and 4G) indicating that counteracting lipid peroxidation could rescue impaired mitochondrial function. It remains to be shown whether this defective mitochondrial function is due to accumulation of phospholipid peroxides in the organelle, as PUFA-phospholipids accumulate there and are beneficial for respiratory growth^69^, or an adaptation mechanism to reduce the production of ROS.

Finally, our work uncovers a previously unnoticed relationship between ferroptosis-protective plasma membrane compartments and quiescence. Despite eisosomes not being conserved in humans, caveolae, which show structural and functional analogies^16^, have recently been reported to be involved in ferroptosis^70–72^. Another interesting analogy is the metabolism of certain cancer stem cells which, similar to quiescent yeasts, rely on oxidative phosphorylation rather than glycolysis^73^. As our results suggest that ubiquinol-dependent protection from ferroptosis is more important in quiescence than fermentation (Figure 4E), it would be interesting to exploit inhibition of ubiquinol-dependent ferroptosis-protective mechanisms as an approach to target quiescent cancer cells, which are resistant to conventional treatments^73, 74^.

## Acknowledgments

We thank T. Zakopoulou and V. Perpiniadis for technical assistance, Dr. S. Sharma, L. Santos Sousa and Dr. G. Kapetanakis for fruitful discussions, S. Limar, S. Walter and S. Schuback for assistance with the lipidomics experiments, Dr. P. Syntichaki for providing materials. This research was co-financed by Greece and the European Union (European Social Fund-ESF) through the Operational Programme «Human Resources Development, Education and Lifelong Learning 2014–2020» in the context of the project “Study of the role of eisosomal proteins in the quiescent state of fungal cells” (MIS 5047827) awarded to CG and VS, and by a grant from the Fondation Santé awarded to CG. FF is a member of the DFG Heisenberg program (FR 3647/4-1) and is supported by the SFB1557. We acknowledge support of this work by the project “The Greek Research Infrastructure for Personalised Medicine (pMED-GR)” (MIS 5002802) which is implemented under the Action “Reinforcement of the Research and Innovation Infrastructure”, funded by the Operational Programme "Competitiveness, Entrepreneurship and Innovation" (NSRF 2014-2020) and co-financed by Greece and the European Union (European Regional Development Fund). A.M. was supported by a doctoral fellowship co-financed by Greece and the European Union (European Social Fund-ESF) through the Operational Programme “Human Resources Development, Education and Lifelong Learning” in the context of the Act “Enhancing Human Resources Research Potential by undertaking a Doctoral Research” Sub-action 2: IKY Scholarship Programme for PhD candidates in the Greek Universities, and by an EMBO Scientific Exchange Grant (9533). We dedicate this manuscript in the memory of Dr. G. Thireos.

## Author contributions

A.M. and C.G. designed the project. A.M. performed and analyzed most of the experiments. A.A. contributed in confocal microscopy and developed scripts for image analysis. D.K. performed and analyzed experiments about ferroptosis. B.E. assisted in the analysis of targeted (oxi)lipidomics. M.M. V.L. M.S. and J.Z. performed measured and analyzed proteomics. M.S. and J.Z. designed proteomics. V.S. provided critical advice on experimental design and data interpretation. B.A. made critical conceptual reorientations of the project. F.F. designed, analyzed, and supervised targeted (oxi)lipidomics and proteomics. C.G. supervised experiments and analyzed data. A.M. and C.G. prepared manuscript. All authors discussed the results, contributed to the manuscript, and agreed on the content of the paper.

## Declaration of interest

The authors declare no competing interests.

## STAR Methods

### Strains, plasmids, and growth conditions

The yeast strains used in this study (Table S7) come from strain Sigma1278b. Only prototrophic strains were used. Cells were grown at 29 °C on a minimal buffered medium pH 6.1^23^, with glucose 3% w/v or lactic acid 0.5 % v/v as the carbon source and ammonium in the form of (NH_4_)_2_SO_4_ (10 mM) as the nitrogen source. Quiescent cells were not formed in non-buffered YNB medium (unpublished results). For liquid cultures, 10^4^ cells/ml were inoculated from precultures in the same medium, for avoiding the lag phase and obtaining high repetitive measurements. Buffered, pH 6.1, YNB was used in the experiments shown in Figure S1E, as Pil1(4A) is not epistatic over the deletion of *NCE102* in our medium^23^. *NCE102-5xGA-GFP* was expressed from a plasmid in a *NCE102*^+^ strain, since the tagged protein could not complement the eisosome disassembly phenotype of the *nce102*Δ strain specifically in quiescent cells (data not shown). For drop tests, cells from OD_600_=8-10 (OD_600_=1-2 in experiments with the *gpx*Δ*flp*Δ), were pelleted, washed in water, diluted to OD_600_=0.3, ternary diluted 3-fold, and 5 μL from the dilutions were spotted on agarose plates. The final concentrations of substances added to solid or liquid media were ethanol 1,18 % w/v, glycerol 2 % v/v, CCCP (30μM), stearic acid (0.1 and 0.3 mM), Oleic acid (0.3 mM, 1 mM, and 6 mM), Linoleic acid (0.1-10 mM), Linolenic acid (0.1-1 mM), α-tocopherol (0.5, 1 mM), bathophenanthrolinedisulfonic acid (BPS) 100 μΜ, unless otherwise stated. Chemicals were purchased from Sigma Aldrich (St. Louis, Missouri, USA) or Cayman Chemical Company (Ann Arbor, Michigan, USA). For survival of the *gpx*Δ*flp*Δ upon linoleic acid (LA) treatment, 10 mM LA was added to exponentially growing cultures at an OD_600_=3, in the presence or not of 1 mM α-tocopherol or 100 μΜ BPS. Knockout, GFP- and mCherry-tagged versions of all genes were created by insertion of cassettes from plasmids, as previously reported^75, 76^, and verified by PCR with external primers and primers specific to the ORF of the respective gene and the selection marker. Plasmids used in this study are listed in Table S7. Primers used are available upon request. *rho0* cells were isolated by treating cells in liquid YPD medium with 10 μg/ml ethidium bromide for 24h at 30 °C, streaking in YPD and testing isolated colonies for growth in YPG (3% v/v Glycerol). *rho0* cells were verified by staining of fixated cells with DAPI (1 μg/ml in PBS) for 5-10 min in the dark and washout with PBS prior to epifluorescence microscopy.

**Cell density measurements** were performed by measuring the OD_600_ of the cultures, properly diluted to fit the linear range of a standard curve, which was calculated from PGD cultures of the *wt* by plotting cell numbers counted in a Neubauer slide with OD_600_ values within the linear range (OD_600_ <0.5) of the spectrophotometer. Statistical significance was assessed using the repeated-measures two-way ANOVA analyses for matched values spread across a row, coupled to Holm-Sidak’s multiple-comparisons test. (Prism, Graphpad, San Diego, CA, USA).

**Glucose measurements** were performed using the GOD/PAP kit (Biotechnological Applications L.T.D., Athens, Greece). Supernatant from cultures was diluted 15-fold in ultrapure water, and 10 μL was mixed with 200 μL reagent in a 96-well plate. Concentrations were normalized to a standard curve and plotted to the OD of the culture. The equation deriving from the scatter plot (Figure S1A) was used to calculate glucose concentrations of the average ODs per timepoint at Figure 1B. **Ethanol measurements** were performed with the ab272531 kit (Abcam PLC, Cambridge, UK), according to the manufacturer’s instructions, for 100 μL of supernatants deproteinated with TCA, from at least 4 independent biological replicate PGD cultures. OD at 580 nm was measured using a Infinite M200 plate reader (Tecan Group Ltd), and the ethanol concentrations were calculated from the standard curve, after subtracting the background of ethanol-depleted culture supernatants. Statistical significance was assessed using the repeated-measures two-way ANOVA analyses for matched values spread across a row, coupled to Holm-Sidak’s multiple-comparisons test. (Prism, Graphpad, San Diego, CA, USA).

**Density gradient separation of cells** was performed two ways, using either Percoll (GE Healthcare, Chicago, Illinois, US)^2^ or Iodixanol (Stemcell technologies, Vancouver, Canada)^77^. Briefly, 1.75 mL of Percoll solution (9V Percoll + 1V NaCl 1.5 M) was centrifuged in 2 mL tubes at 19,320 g, 20 °C with an angle rotor for 15 min to form continuous densities gradient (1.00-1.31 g/mL). 100 μL sample (2 ml form PGD cultures, precipitated, washed, and resuspended in 1 M Tris pH 7.4) was carefully overlaid onto the preformed continuous density gradient, and centrifuged at 400 g for 60 min in an angle rotor tabletop centrifuge at 20 °C. For iodixanol, 500 μL of cells were laid on top of 500 μL of a 26 % solution in 10 mM Tris-HCl pH 7.5, 150 mM NaCl, in 2 mL tubes. Tubes were centrifuged at 2500 g for 2.5 min in an angle rotor tabletop centrifuge at 20 °C, rotated by 180 degrees and re-centrifugated. The fractions were collected, washed once with 500 μL Tris buffer (pH = 7.4), pelleted and resuspended in Tris buffer or conditioned medium for subsequent assays or sterile water.

### Confocal microscopy, quantifications, and statistics

Images in Belgium were acquired on a Zeiss LSM710 microscope equipped with a 100× differential interference contrast, numerical aperture (NA) 1.46 Alpha-Plan-Apochromat objective, Airyscan module and ZEN 2.1 SP2 software. Airyscan processing was done using the default settings of the ZEN software. The final pixel size was 38.5 nm. Images in Athens were acquired on a Leica TCS SP8 MP with AOBS (acousto optical beam splitter), equipped with HC Plan APO 63x, NA 1.40 oil immersion objective. The final pixel size was 25.8 nm. GFP and mCherry fluorescence were excited with the 488-nm line of the argon laser and a 594 nm solid-state laser, respectively. Appropriate filters and beam splitters were used. In all conditions, gain percentage and laser intensities were kept to levels of non-saturated signal acquisition. The same laser intensities and other settings of the confocal microscope were used for all image acquisition in a single timepoint. For comparison between timepoints with important differences in signal, only laser intensity or gain were changed, and the ratios of the change were considered during quantifications. Images were smoothed, corrected for optimal brightness and contrast, cropped and merged using FIJI software (National Institutes of Health, Bethesda, MD, USA)^78^, and annotated with Inkscape (https://inkscape.org). In each figure only a few cells representative of the whole cell population, observed in at least one preliminary experiment on epifluorescence microscopy and two independent biological replicate experiments on confocal, are shown.

For quantifications, images were analyzed with custom-made FIJI macros, calculating the normalized mean fluorescence intensity, the MCC-to-total surface area or total-to-internal fluorescence intensity, as previously^23, 79^. Briefly, for total-to-internal fluorescence intensity, two homocentric ellipses outlining the whole cell or the whole cells excluding the plasma membrane were manually drawn in middle-section images. For the other parameters, a mask was first created in the image of the MCC marker, in surface-section images. A 3×3 median filter (despeckle function) was applied, the ImageJ default thresholding method was applied, and then overlapping MCCs were separated using the watershed function. The intensities of the channels of interest were measured within manually selected cell outlines, while the median of the fluorescence intensity in the whole image was subtracted as background. The MCC-to-total surface area was defined as the ratio of the area of MCC-positive objects to the area of the outlined region. The Pearson’s correlation coefficient was obtained using a custom-made macro based on the coloc2 plugin of FIJI. Due to the much higher number of MCCs at PGD timepoints, our custom-made script sometimes tended to recognize closely localized MCCs as one, leading to a slight overestimation of the MCC/total surface ratio. All parameters were calculated from at least two independent biological replicates for each condition. The values for single cells are presented in box-and-whisker plots. After verification that two independent biological replicates gave statistically nonsignificant differences in mean values, the values of the two experiments were merged. Prism software (Graphpad, San Diego, CA, USA), one-way ANOVA with the nonparametric Kruskal–Wallis test, and Dunn’s multiple-comparison post hoc analyses were used to assess the significance of the value differences of all measurements.

### Determination of the relationship between eisosome induction and the morphology of mitochondria

4 independent biological replicate cultures of a Pil1-mCherry- and Ilv3-GFP-expressing strain were imaged by confocal microscopy at different timepoints PGD (n total = 5028). Cells were qualitatively sorted in different categories, by 2 experienced investigators, regarding the mitochondrial morphology and the eisosome expansion/assembly. For mitochondrial morphology the distribution of Ilv3-GFP was used and the three following categories were considered, according to Laporte et al. 2018^6^: 1) cells with no green fluorescence, reportedly dead 2) cells with only 4 or less aggregates of green fluorescence, which were considered senescent, and 3) the rest of the cells which displayed several vesicles, which were considered possessing cortical mitochondria and thus, quiescent. Regarding eisosome expansion, cells were first manually sorted in two categories: 1) cells with induced Pil1, and 2) cells without Pil1 induction. The validity of this manual classification was verified by measuring Pil1 intensity in representative cells from selected timepoints of 1 biological replicate (Figure S2C). An additional criterion of classification about eisosome assembly was the topology of Pil1-mCherry, which was either mainly at the periphery of the cells, or cytoplasmic or an intermediate situation. This eventually led us to adopt a total of 6 categories of eisosomal states, summarized in Figure 2A and for which representative images are shown in Figure 2B. Results in Figure 2C show percentages of these cell categories at different timepoints PGD, from 4 biological replicates. The physiological significance of all these categories was not further investigated, and only the main categories regarding induction of Pil1 or not have been analyzed further. Cell numbers in all categories were summed (Table S2) and the probability of a cell with or without eisosome induction to be Quiescent, senescent, or dead (Table S3) was calculated and expressed as pie charts. Chi-squared tests for each timepoint were performed to determine the statistical significance of independence of the values. Chi-squared values and p-values for each time point are shown in Table S2.

**Epifluorescence microscopy** images were obtained with an Axioplan 2 (Carl Zeiss, Inc) microscope equipped with Plan-Apochromat x100 1.40 NA oil immersion objective lens, and appropriate fluorescence light filter sets. Images were captured with a Retiga3 (QImaging, Surrey, BC, Canada) digital camera and Occular 2.01 acquisition software (Teledyne Photometrics, Tucson, AZ, USA). Images were processed with FIJI^78^ and annotated with Inkscape (https://inkscape.org). Staining of dead cells with methylene blue was performed by mixing an equal volume of cell culture with a 0.1% methylene blue, 2% sodium citrate solution. For visualization of stress granules, cells were incubated at 37 °C for 30 min, as previously^27^, and manually measured from 2 independent experiments. Cells from liquid cultures were laid on a thin layer of 1% agarose supplemented with appropriate nutrients and observed at room temperature. In each figure, a few representative cells, observed in at least two independent biological replicate experiments, are shown.

### Western blotting

Total cell protein extracts were prepared and analyzed by SDS–PAGE as previously^79^. Proteins were transferred to a PVDF membrane (Immobilon, Macherey Nagel, Düren, NW, Germany) and probed with a mouse monoclonal anti-GFP (Roche, Basel, Switzerland), mouse monoclonal anti-mCherry (ab125096, Abcam PLC, Cambridge, UK) or anti-actin (sc47778, Santa Cruz Biotechnology, Dallas, TX, USA). Primary antibodies were detected by enhanced chemiluminescence (Millipore, Burlington, MA, USA) after treatment with horseradish-peroxidase-conjugated anti-mouse immunoglobulin (Ig) G secondary antibody (Cell Signalling, Danvers, MA, USA). Signals were detected with ImageQuant LAS 4000 mini (FujiFilm, Tokio, Japan), and images were corrected for optimal brightness and contrast with FIJI and annotated with Inkscape. Relative semi-quantitative amounts of total proteins were estimated from 1–4 biological replicates of non-saturated exposures using the gel analyzer tool of FIJI^78^. Each band was selected by using rectangular ROI selection and “Gels” analyzer, followed by quantification of the peak area of obtained histograms. Data were acquired as area values. In each graph, the ratios of signal/actin are plotted in scatter dot plots, normalized to the ratio of EXP which is set as 1. Statistical significance was assessed using the repeated-measures two-way ANOVA analyses for matched values spread across a row, coupled to Holm-Sidak’s multiple-comparisons test. (Prism, Graphpad, San Diego, CA, USA).

### Estimation of cell survival

Cells from cultures at different timepoints were normalized to OD_600_=0.9, diluted decimally 3-fold, and 30 μL were spread on YPD plates, incubated for 2-3 days at 30 °C and Colony Forming Units (CFUs) were counted. 2 technical replicates were performed for each condition and averaged. CFUs are expressed as percentage of total cells in the sample, which was estimated by cell density measurements. The number of total cells calculated were verified by CFU counting of *wt* cells from the GD timepoint, considering that at this stage 10% of the cells are senescent or dead (Figure 2C). Independent biological replicates are shown in box and whiskers plots. Unpaired two tailed students T-tests at indicated timepoints were used to calculate the significance of the value differences.

### Proteomics

For determination of the proteome of EXP vs PGD cells, 3 biological replicates of cultures of the *wt* strain in minimal buffered medium were inoculated, and samples of at least 10^8^ cells were collected at an OD_600_=1, and 30 h later (OD_600_=18), washed three times with PBS and stored at -80 °C. Samples were homogenized in FASP lysis buffer (4% SDS, 0.1M DTE, 0.1M Tris-HCl pH 7.6). Protein concentration was determined by Bradford assay. Protease inhibitors (Roche, Basel, Switzerland) were added at a final concentration of 3.6% and samples were stored at -80°C until further use. Protein extracts (200 μg/sample) were processed using filter aided sample preparation (FASP) as described previously^80^, with minor modifications^81^. Briefly, buffer exchange was performed in Amicon Ultra Centrifugal filter devices (0.5 mL, 30 kDa MWCO; Merck) at 14,000 rcf for 15 min at room temperature. The protein extract was mixed with urea buffer (8M urea in 0.1M Tris-HCl pH 8.5) and centrifuged. The concentrate was diluted with urea buffer and centrifugation was repeated. Alkylation of proteins was performed with 0.05M iodoacetamide in urea buffer for 20 min in the dark followed by a centrifugation at 14,000 rcf for 10 min at RT. Additional series of washes were conducted with urea buffer (2 times) and ammonium bicarbonate buffer (50 mM NH_4_HCO_3_ pH 8.5, 2 times). Tryptic digestion was performed overnight at RT in the dark, using a trypsin to protein ratio of 1:100. Peptides were eluted by centrifugation at 14000 rcf for 10 min, lyophilized and stored at –80°C until further use. The peptides were purified using a modified Sp3 clean up protocol and finally solubilized in the mobile phase A (0.1% Formic acid in water), sonicated and the peptide concentration was determined through absorbance at 280nm measurement using a nanodrop instrument. Samples were analyzed on a liquid chromatography tandem mass spectrometry (LC-MS/MS) setup consisting of a Dionex Ultimate 3000 nanoRSLC coupled in line with a Thermo Q Exactive HF-X Orbitrap mass spectrometer. Peptidic samples were directly injected and separated on an 25 cm-long analytical C18 column (PepSep, 1.9μm3 beads, 75 µm ID) using an one-hour long run, starting with a gradient of 7% Buffer B (0.1% Formic acid in 80% Acetonitrile) to 35% for 40 min and followed by an increase to 45% in 5 min and a second increase to 99% in 0.5min and then kept constant for equilibration for 14.5min. A full MS was acquired in profile mode using a Q Exactive HF-X Hybrid Quadrupole-Orbitrap mass spectrometer, operating in the scan range of 375-1400 m/z using 120K resolving power with an AGC of 3x 106 and maximum IT of 60ms followed by data independent acquisition method using 8 Th windows (a total of 39 loop counts) each with 15K resolving power with an AGC of 3x 105 and max IT of 22ms and normalized collision energy (NCE) of 26. Each sample (e.g. three biological replicas) was analyzed in three technical replicas. Orbitrap raw data were analyzed in DIA-NN 1.8.1 (Data-Independent Acquisition by Neural Networks) through searching against the *Saccharomyces cerevisiae* Reference Proteome (downloaded from Uniprot, 6052 proteins entries, downloaded 30/5/2022) using the library free mode of the software, allowing up to two tryptic missed cleavages and a maximum of three variable modifications/peptide. A spectral library was created from the DIA runs and used to reanalyse them (double search mode). DIA-NN search was used with oxidation of methionine residues and acetylation of the protein N-termini set as variable modifications and carbamidomethylation of cysteine residues as fixed modification. The match between runs feature was used for all analyses and the output (precursor) was filtered at 0.01 FDR and finally the protein inference was performed on the level of genes using only proteotypic peptides. The proteomics data were processed in Perseus 1.6.15.0^82^. Values were log(2) transformed, a threshold of 70% of valid values in at least one group was applied and the missing values were replaced from normal distribution. For statistical analysis, Student’s t-test was performed and permutation-based FDR was calculated. The mass spectrometry proteomics data have been deposited to the ProteomeXchange Consortium via the PRIDE^83^ partner repository with the dataset identifier PXD041191.

For determining the changes in the proteomes of *mcc*Δ and *flp*Δ mutants compared to *wt* cells, label free proteomics were performed. Triplicates of 2 OD units of *wt* and *mcc*Δ cells were collected by centrifugation, 2 days PGD. The pellets were resuspended in lysis buffer of the commercially available proteomics kit (iST Kit, preomics) and peptides were prepared according to the manufacturer’s recommendations. A mixture of trypsin and LysC was used to perform protein digestion. Reversed-phase chromatography was performed using a Thermo Ultimate 3000 RSLCnano system coupled with a Q ExactivePlus mass spectrometer (Thermo) via a nano-electrospray ion source, as previously described with some adjustments. Peptides were eluted from a 50 cm C30 column using a gradient of acetonitrile from 1-12 % for 45 min and from 12-35% in 0.1% formic acid for 70 min, followed by 35-60% for 20 min and 60-90% for 10 min at a constant flow rate of 200 nl/min. The 10 most intense multiply charged ions (z = 2-5) from the survey scan were selected with an isolation width of 1.4 m/z. Sequenced peptides were dynamically excluded for 20 s. The resulting spectra were analyzed with MaxQuant (version 2.1.4.0, www.maxquant.org; Cox and Mann, 2008; Cox et al., 2011). Label free analysis was performed with Perseus^82^. The mass spectrometry proteomics data have been deposited to the ProteomeXchange Consortium via the PRIDE^83^ partner repository with the dataset identifier PXD041123 and 10.6019/PXD041123.

Volcano-plots were created using VolcaNoseR2 online tool^84^. Each plot summarizes only the significant differences [log_2_(fold-change)] in protein abundance in the x-axis and the –log_10_(p-value) of the Student’s t-test performed from replicates in the y-axis. The colored dots correspond to upregulated (purple) or downregulated (blue) proteins and the gray to the unaffected proteins based on the cut-off set in each plot. For Figure 1C, the cut-off of difference was set at 1.9 and the significance of Student’s t-test at p<0.05. For Figures 3F, and S3G, the cut-off of difference was set at 0.5 and the significance of Student’s t-test at p<0.05.

GO-term enrichment for biological processes and cellular components, for proteins with Student’s T-test p-value <0.05 and the indicated difference, was performed with PANTHER^85^ using the default sorting for fold-enrichment and hierarchical clustering of the overrepresented functional classes. In Figure 1D, S1B and 3G only the most specific subclasses are shown. Mean fold-changes of the protein constituents of the yeast cellular components in Figure 1D were calculated in Microsoft Excel (using the SUMPRODUCT equation), for all the proteins of the dataset with T-test p-value <0.05. The proteins of the indicated yeast GO-Slim cellular components in Figure 1D were acquired from the *Saccharomyces* Genome Database.

### Fatty acid supplementation and targeted lipidomics for Phosphatidylethanolamine

For LC-MS/MS analysis of Phosphatidylethanolamine (PE) species, cells were grown at 29 °C on a minimal buffered medium, pH 6.1^23^ with glucose 3% as carbon source and ammonium as nitrogen source. Cultures were supplemented or not with the indicated concentrations of Fatty acids. Fatty acids were purchased from Sigma [Oleic acid (O1383), Linoleic acid (L1012), Linolenic acid (L2376)]. Cell pellets (19 OD units) were resuspended in 0.75 ml of 150 mM ammonium formate and lysed at 4 °C with 500 μl of acid-washed glass beads using FastPrep Homogenizer (MP Biomedicals). The protein concentration of each sample was determined using the Bradford reagent (Bio-Rad). Lipids were extracted from lysed yeast cells according to 50 μg of protein by chloroform/methanol extraction^86^. Prior to extraction, ceramide (17:0/18:1 Avanti Polar Lipids) was spiked into each sample for normalization. Dried lipids were dissolved in a 65:35 mixture of mobile phase A (50:50 water/acetonitrile, including 10 mM ammonium formate and 0.1% formic acid) and mobile phase B (88:10:2 2-propanol/acetonitrile/H20, including 2 mM ammonium formate and 0.02% formic acid). HPLC analysis was performed on a C30 reverse-phase column [Accucore C30 LC column (150 mm x 2.1 mm 2.6 µm Solid Core; Thermo Fisher Scientific)] connected to aNexera XR HPLC (Shimadzu). A binary solvent system (mobile phases A and B) was used in a 6-min gradient at a flow rate of 0.4 ml/min and an injection volume of 1 μl. For the gradient 40% B for 0.1 min was used followed by its increase from 40% to 50% over 2 min. Afterwards, Buffer B was increased from 50% to 100% over 1 min. 100% B was kept for 1 min and decreased to 40 % B for 0.1 min. 40% B was kept until the end of the gradient. Lipid samples were analyzed by a Qtrap 5500 (Thermo Fisher Scientific) equipped with a heated electrospray ionization (HESI) probe. The MS was operated in negative ion mode using multiple reaction monitoring mode with 60-s detection windows. The predefined transitions for the PE species are provided in Table S6. Peaks were analyzed using the SciexOS software as described previously^87^. Peaks were defined through raw files, product ion and precursor ion accurate masses. From the intensities of the lipid standard (PE 16:0/18:1, Avanti Polar Lipids), absolute values for each lipid in pmol/μg protein were calculated. Data from 2-6 replicates are displayed as a percentage of each phospholipid PE species normalized to the total PE detected in each sample. The lipidomics workflow was reported in the Lipidomics Standards Initiative (LSI) platform (https://lipidomicstandards.org/). The report details are provided in Table S8.

## Supplemental Information titles and legends

**Figure S1. Extended data of Figure 1.**
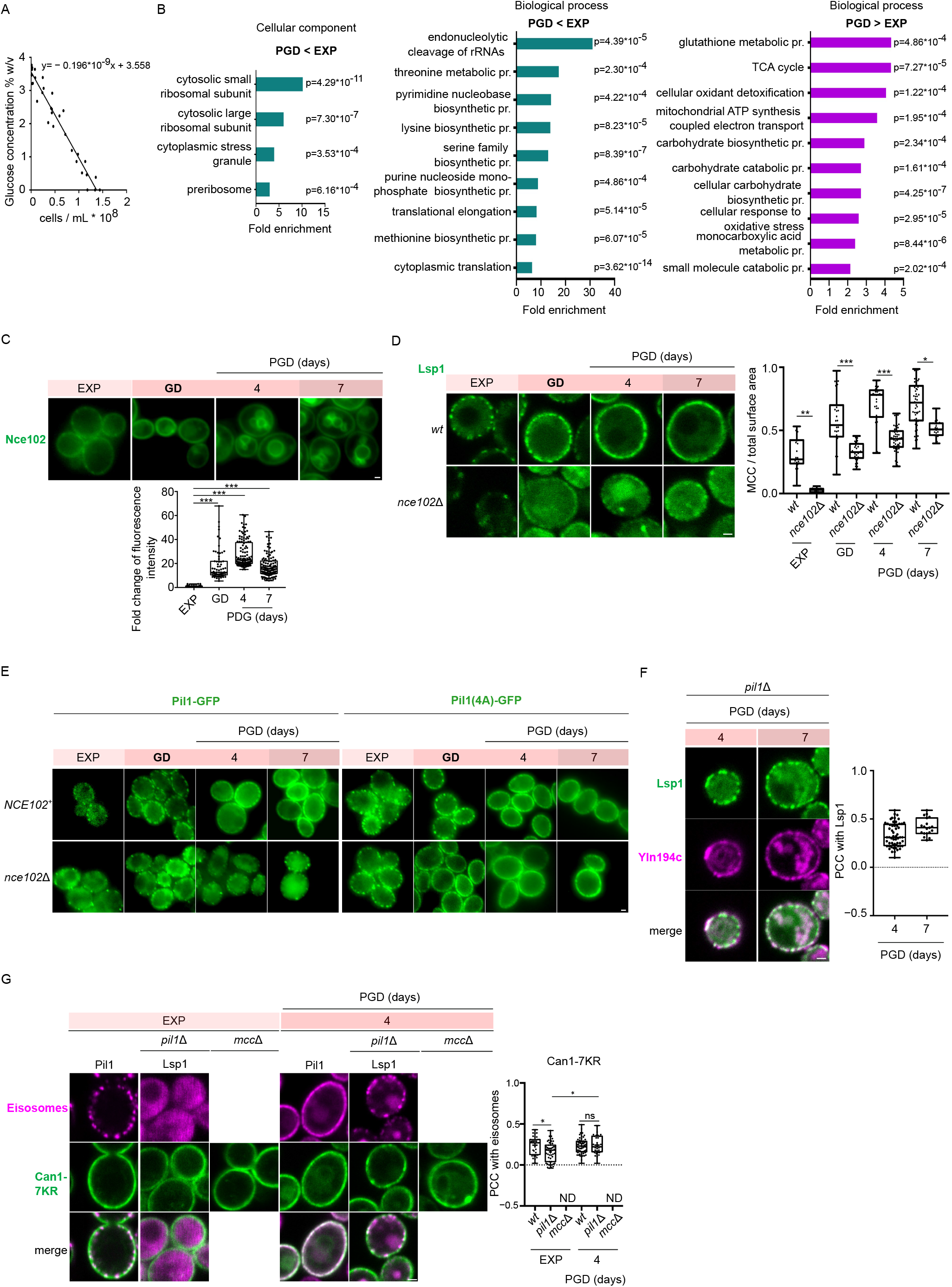
Eisosomes expand upon glucose depletion, and this requires the BAR-domain Lsp1 protein. **A)** Scatter plot of glucose concentration vs optical density in supernatants of yeast cultures. The equation derived from the non-linear regression of fit analysis of the data from 3 independent experiments is shown. **B)** Bar graphs of the top GO-TERM enriched cellular components and biological processes in downregulated (green), and upregulated (magenta) proteins of Figure 1C. The fold enrichment and p-values are shown. **C)** Middle section epifluorescence microscopy of a Nce102-GFP-expressing *NCE102^+^* strain, at EXP and different timepoints PGD. Quantifications: Nce102-GFP single-cell fluorescence intensities (n = 67-121 cells), normalized to the signal from EXP cells, are plotted in box and whiskers plots. \*\*\**P* < 0.001, from one-way ANOVA. **D)** Middle section confocal microscopy of Lsp1-GFP-expressing *wt* and *nce102*Δ strains, at EXP and different timepoints PGD. Quantifications: The single-cell MCC/Total surface area ratios from surface section images (n=20-84), are plotted in box and whiskers plots. \*\*\**P* < 0.001. \**P* < 0.05; ns, non-significant, p > 0.05, from one-way ANOVA. **E)** Epifluorescence microscopy of Pil1-GFP- or Pil1(4A)-GFP-expressing *wt* and *nce102*Δ strains, at EXP or different timepoints PGD. As in EXP^18^, the *PIL1(4A)* is epistatic over the deletion of *NCE102*, indicating that Nce102 promotes eisosome assembly via inhibition of Pil1 phosphorylation by Pkh1/2^88^. **F)** Middle section confocal microscopy of a Lsp1-GFP- and Ynl194c-mCherry-expressing *pil1*Δ strain, at different timepoints PGD. Quantifications: The Pearson correlation coefficient (PCC) of the two fluorescence signals from single cells (n=19-57) from surface-section images are plotted in box and whiskers plots. **G)** Middle section confocal microscopy of Can1(7KR)-GFP and Pil1-mCherry or Lsp1-mCherry-expressing *wt, pil1*Δ and *pil1*Δ*lsp1*Δ (*mcc*Δ) strains, at EXP and 4 days PGD. Quantifications: The Pearson correlation coefficient (PCC) of the two fluorescence signals in single cells (n=35–60) from surface-section images are plotted in box and whiskers plots. N.D. not defined \**P* < 0.05; ns, non-significant, p > 0.05, from one-way ANOVA. Can1 in *mcc*Δ is much more homogeneous at the plasma membrane of both EXP and PGD cells. Only a few eisosome remnants are seen, as previously described for EXP^12^. (Scale bar: 1 μm).

**Figure S2. Extended data on Figure 2.**
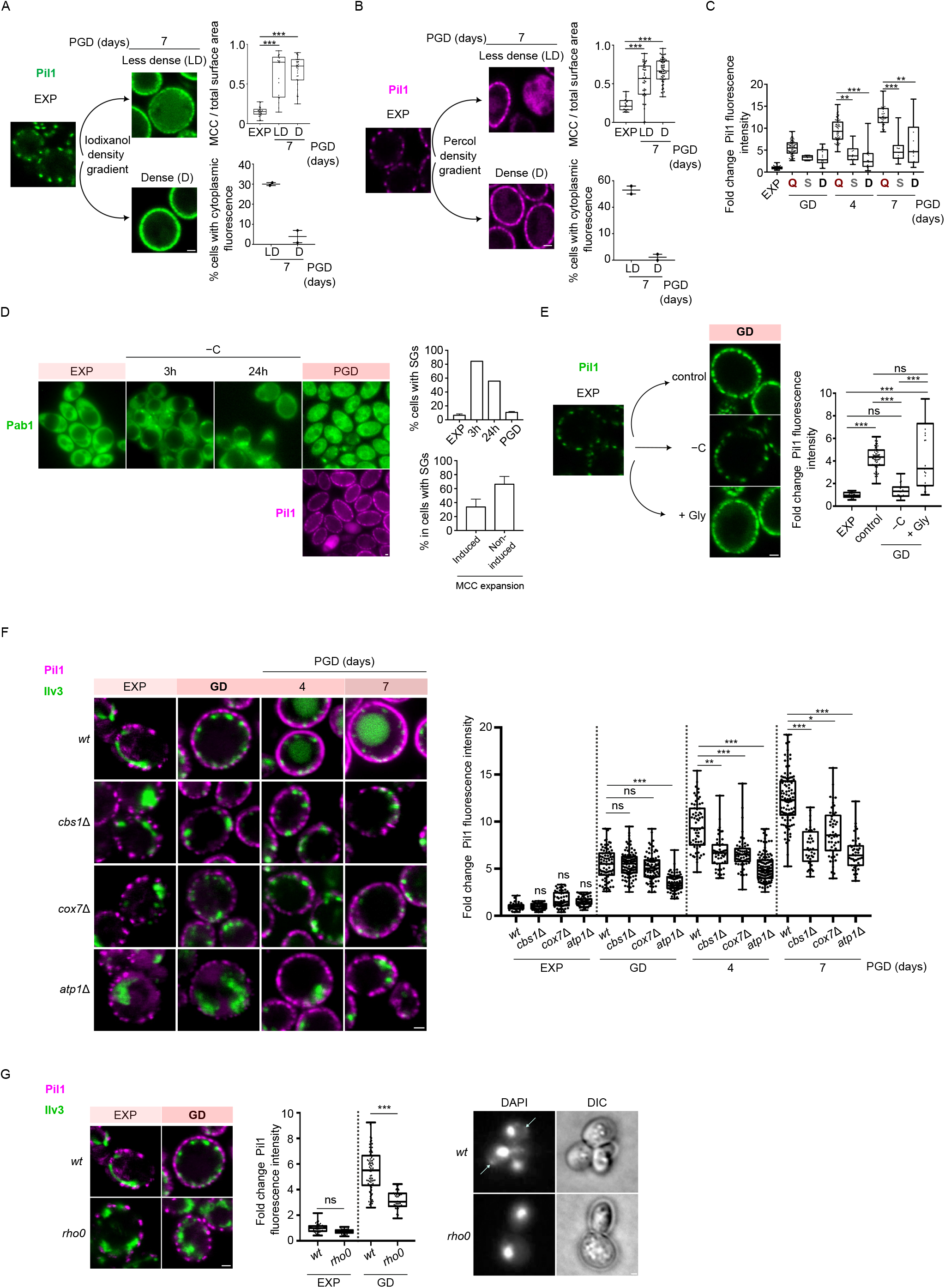
Eisosomes expand in respiratory-active quiescent yeasts. **A), B)** Middle section confocal microscopy of a Pil1-GFP- or Pil1-mCherry-expressing strain, from EXP and 7 days PGD. The latter samples are separated with iodixanol (A) or percoll (B) density-gradient centrifugation into dense (D) and less dense (LD) cells. Quantifications: The single-cell MCC/Total surface area (n=17-58) from surface-section images and the percentage of cells with cytoplasmic Pil1 (n=44-239) are plotted in box and whiskers or scatter plots (mean±s.d), respectively. \*\*\**P* < 0.001, from one-way ANOVA. **C)** Pil1-mCherry single-cell fluorescence intensities from representative quiescent (Q), senescent (S) and dead (D) cells (n=4-74) at the indicated timepoints of the experiment in Figure 2C, normalized to the signal from EXP cells. Data from 1 representative (out of four) experiments are plotted in box and whiskers plots. \*\*\**P* < 0.001; \*\**P* < 0.01, from one-way ANOVA. **D)** Stress granules do not form PGD, contrary to carbon starvation. Epifluorescence microscopy images of a strain expressing Pab1-GFP and Pil1-mCherry, at EXP, PGD, or shifted from EXP to carbon starvation (–C) for 3 or 24 h. Quantifications: The percentage of cells (n=97-267) displaying stress granules at different conditions, and, within the stress granule-possessing cells (n=43) at PGD timepoint the percentage of cells with induction of Pil1 or defective eisosome assembly/induction, from two independent experiments are plotted in bar charts as mean ± s.d. **E)** Middle section confocal microscopy of a Pil1-GFP-expressing strain, at EXP or PGD (control), or shifted to carbon-free medium, supplemented with 2% w/v Glycerol (+Gly) or not (–C). Quantifications: Pil1-GFP single-cell fluorescence intensities (n=23–68), normalized to the signal from EXP cells, are plotted in box and whiskers plots. \*\*\**P* < 0.001; ns, non-significant, p > 0.05, from one-way ANOVA. **F)** Middle section confocal microscopy of Pil1-mCherry- and Ilv3-GFP-expressing *wt* or respiration-deficient strains, at different timepoints PGD, Quantifications: Pil1-mCherry single-cell fluorescence intensities (n=36-118), normalized to the signal from EXP *wt* cells, are plotted in box and whiskers plots. \*\*\**P* < 0.001; \*\**P* < 0.01; \**P* < 0.05; ns, non-significant, p > 0.05, from one-way ANOVA. **G)** Left: Middle section confocal microscopy of wt and *rho0* Pil1-mCherry- and Ilv3-GFP-expressing strains, at EXP or at GD, Quantifications: Pil1-mCherry single-cell fluorescence intensities (n=32-74), normalized to the signal from EXP *wt* cells, are plotted in box and whiskers plots. \*\*\**P* < 0.001; ns, non-significant, p > 0.05, from one-way ANOVA. Right: epifluorescence microscopy of DAPI-staining of the *wt* and *rho0* strains, confirms the absence of mitochondrial DNA in the latter. Arrows point to the mitochondrial staining of DAPI in *wt*. (Scale bar: 1 μm).

**Figure S3. Extended data of Figure 3.**
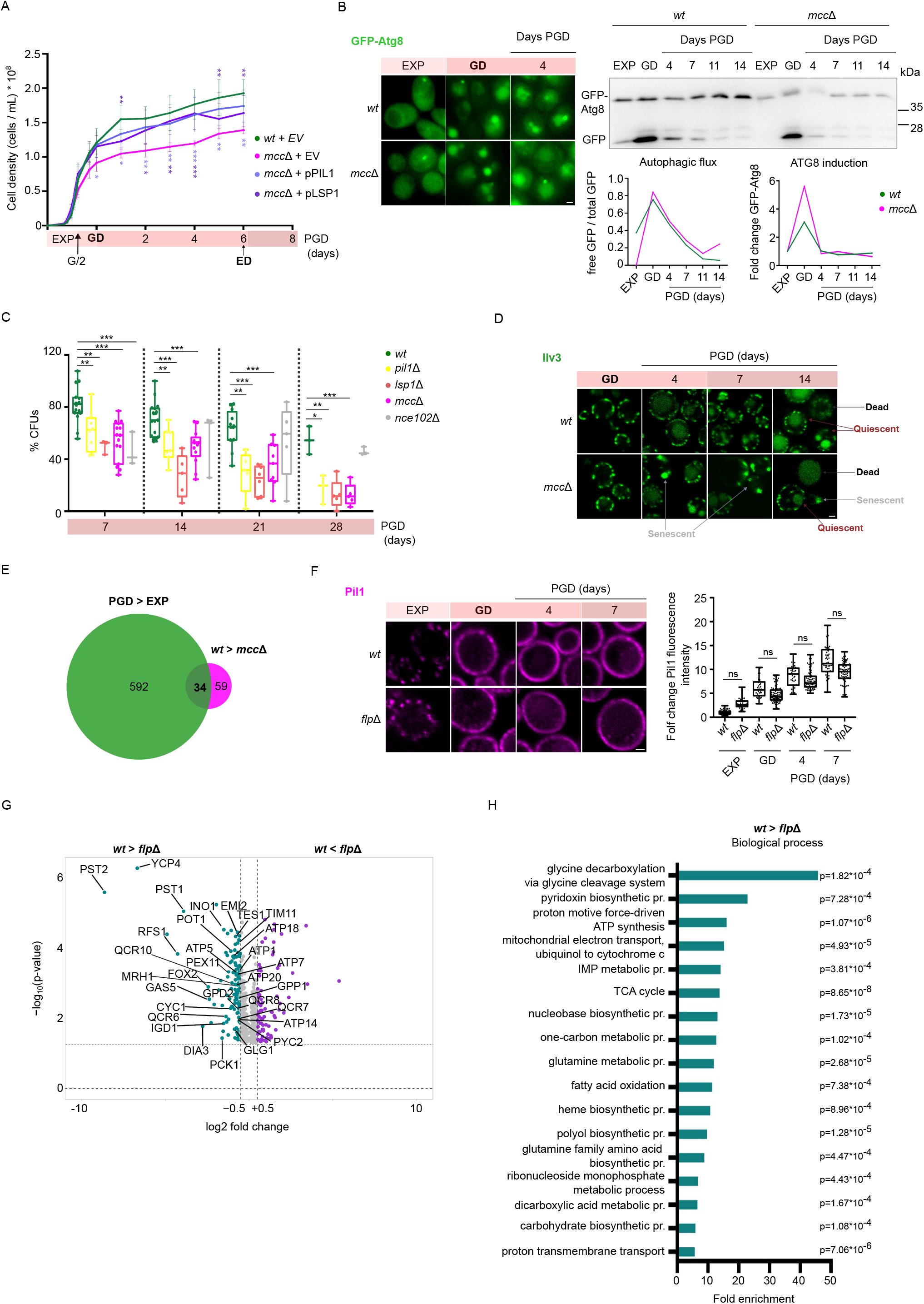
Eisosomes promote respiratory growth and are required for the long-term quiescence maintenance by stabilizing the Flavodoxin-Like Proteins. **A)** Line chart of *wt*, and *pil1*Δ*lsp1*Δ (*mcc*Δ) strains transformed with either empty vector (EV) or plasmid-borne *PIL1* (pPIL1) or *LSP1-GFP* (pLSP1), cultivated in minimal medium. Cell density measurements at different timepoints in EXP and PGD, from 5-6 independent experiments are plotted as mean ± s.d. \*\*\**P* < 0.001; \*\**P* < 0.01; \**P* < 0.05, p > 0.05, from two-way ANOVA. **B)** The *mcc*Δ strain has normal induction of autophagy PGD. Left: Epifluorescence microscopy images of *wt* and *mcc*Δ strains transformed with a GFP-ATG8-expressing plasmid, from EXP, GD and 4 days PGD. Right: Western blot of total protein extracts from the same strains, collected at EXP or different timepoints PGD, probed with anti-GFP. The 2 critical components of the autophagic response where quantified: 1) for the autophagic flux the ratio of the signal intensities of the free GFP to total GFP is plotted and 2) for the *ATG8* induction the signal intensities of GFP-Atg8 were normalized to EXP of each strain were plotted. **C)** Growth recovery of *wt*, *pil1*Δ, *lsp1*Δ, *mcc*Δ, and *nce102*Δ yeast cultures in minimal medium, as percentages of CFUs, from 3-19 independent experiments, are plotted in box and whiskers plots. \*\*\**P* < 0.001; \*\**P* < 0.01; \**P* < 0.05, from unpaired T-tests. **D)** Middle section confocal microscopy images of Ilv3-GFP-expressing *wt* and *mcc*Δ strains at various timepoints PGD. Quiescent (vesicular-mitochondria), Senescent (globular mitochondria) and Dead (no Ilv3-GFP fluorescence, only vacuolar autofluorescence) are indicated. **E)** Venn diagram of the proteins downregulated in *mcc*Δ vs *wt* at 3 days PGD (Figure 3F) and the proteins upregulated in *wt* upon GD (Figure 1C). **F)** Middle section confocal microscopy of Pil1-mCherry-expressing *wt* and *pst2*Δ*ycp4*Δ*rfs1*Δ (*flp*Δ) strains, at EXP and different timepoints PGD. Pil1-mCherry single-cell fluorescence intensities (n=35-90), normalized to the signal from *wt* EXP cells, are plotted in box and whiskers plots. \*\*\**P* < 0.001. \**P* < 0.05; ns, non-significant, p > 0.05, from one-way ANOVA (Scale bar: 1 μm). **G)** Volcano-plot [Student’s T-difference vs −log_10_(p-value)] comparison of the proteome of *pst2*Δ*ycp4*Δ*rfs1*Δ (*flp*Δ) vs *wt* cells at 3 days PGD. Proteins with significant >0.50 (magenta) and <–0.50 (green) log_2_(fold-change) are indicated. FLPs, proteins downregulated in *mcc*Δ, mitochondrial proteins, proteins involved in fatty acid oxidation and carbohydrate metabolism are annotated. **H)** bar graph of the top GO-TERM enriched biological processes in downregulated proteins of (G). The fold enrichment and p-values are shown.

**Figure S4. Extended data of Figure 4.**
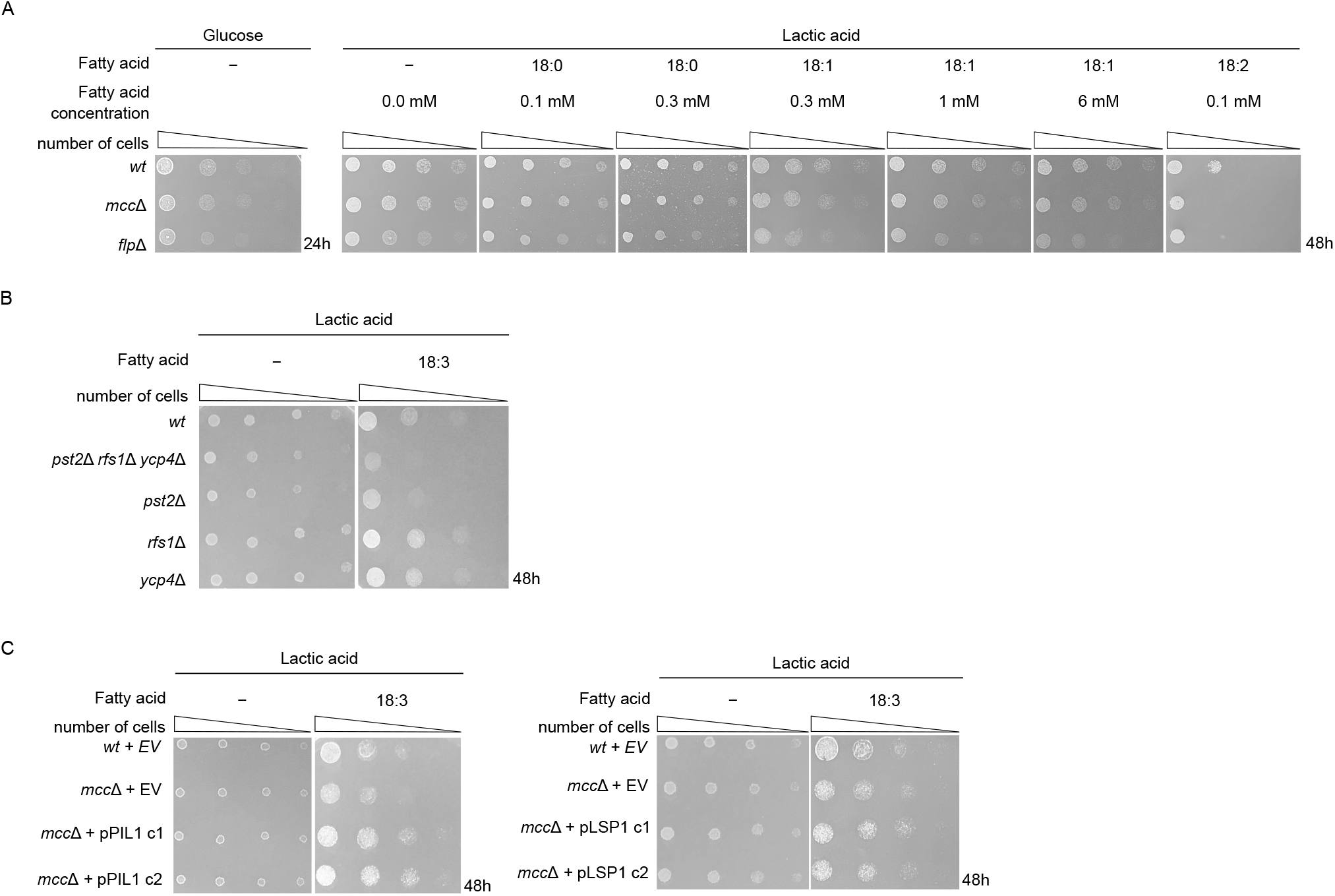
Eisosomes and Flavodoxin-Like Proteins protect Quiescent cells from lipid peroxidation and ferroptosis. **A)** Drop tests of *wt*, *pil1*Δ*lsp1*Δ (*mcc*Δ) and *pst2*Δ*ycp4*Δ*rfs1*Δ (*flp*Δ) strains in minimal medium with glucose or lactic acid as carbon source, supplemented or not with the indicated concentrations of the indicated fatty acids. **B)** Drop tests of *wt*, *flp*Δ, *pst2*Δ, *ycp4*Δ, and *rfs1*Δ strains in minimal medium with lactic acid as carbon source, supplemented or not with 0.1 mM Linolenic acid (LNA). **C)** Drop tests of *wt* and *mcc*Δ strains transformed with either empty vector (EV) or plasmid-borne *PIL1* (pPIL1) or *LSP1-GFP* (pLSP1) [two different colonies (c1, c2)] in minimal medium with lactic acid as carbon source, supplemented or not with 0.1 mM Linolenic acid (LNA).

## Supplemental tables

Table S1: Statistically processed mass spectrometry data sets, related to Figures 1C, 1D, S1B.

Table S2: Cell numbers and statistical analysis for the correlation of mitochondrial distribution and the eisosome-induction/assembly PGD.

Table S3: Probabilities of eisosome-expansion and/or assembly as predictive markers of Quiescence.

Table S4: Statistically processed mass spectrometry data sets, related to Figures 3F, 3G, S3H and S3G.

Table S5: Statistical analysis of Figure 4F. Holm-Sidak’s multiple comparisons test.

Table S6: List of transitions used for targeted lipidomics. Table S7. *S. cerevisiae* strains and plasmids used in this study.

## Notes

### Competing Interest Statement

The authors have declared no competing interest.

