## Supplementary material for "Ferroptosis-Protective Membrane Domains in Quiescence": Tables S2_S3_S5_S6_S7

**Table S2: Cell numbers and statistical analysis for the correlation of mitochondrial distribution and the eisosome-induction/assembly PGD.**

|  |  | **Induced Pil1** | | **Non-induced Pil1** | | | | |  |  |
| --- | --- | --- | --- | --- | --- | --- | --- | --- | --- | --- |
| **days PGD** | mitochondrial morphology | Induced Pil1_Many eisosomes | Induced Pil1_slightly cytoplasmic pil1 | Few eisosomes_membrane Pil1 | Few eisosomes_cyt Pil1 | Cyt Pil1 | No Pil1 signal | total | Chi-squared | p-value |
| **0** | Vesicular | 536 | | 33 | | | | 569 | 310.1, 2 | <0,0001 |
|  |  | 462 | 74 | 0 | 23 | 10 | 0 |  |  |  |
|  | Globular | 9 | | 34 | | | | 43 |  |  |
|  |  | 6 | 3 | 4 | 10 | 20 | 0 |  |  |  |
|  | Dead | 1 | | 22 | | | | 23 |  |  |
|  |  | 0 | 1 | 0 | 3 | 19 | 0 |  |  |  |
|  | total | 546 | | 89 | | | | 635 |  |  |
|  |  | 468 | 78 | 4 | 36 | 49 | 0 |  |  |  |
| **4** | Vesicular | 837 | | 24 | | | | 861 | 574.8, 2 | <0,0001 |
|  |  | 831 | 6 | 10 | 10 | 4 | 0 |  |  |  |
|  | Globular | 18 | | 63 | | | | 81 |  |  |
|  |  | 18 | 0 | 16 | 9 | 36 | 2 |  |  |  |
|  | Dead | 19 | | 50 | | | | 69 |  |  |
|  |  | 19 | 0 | 0 | 4 | 27 | 19 |  |  |  |
|  | total | 874 | | 137 | | | | 1011 |  |  |
|  |  | 868 | 6 | 26 | 23 | 67 | 21 |  |  |  |
| **7** | Vesicular | 1493 | | 10 | | | | 1503 | 1365, 2 | <0,0001 |
|  |  | 1488 | 5 | 5 | 2 | 3 | 0 |  |  |  |
|  | Globular | 8 | | 86 | | | | 94 |  |  |
|  |  | 8 | 0 | 57 | 14 | 14 | 1 |  |  |  |
|  | Dead | 6 | | 47 | | | | 53 |  |  |
|  |  | 6 | 0 | 3 | 0 | 25 | 19 |  |  |  |
|  | total | 1507 | | 143 | | | | 1650 |  |  |
|  |  | 1502 | 5 | 65 | 16 | 42 | 20 |  |  |  |
| **11** | Vesicular | 596 | | 0 | | | | 596 | 348.5, 2 | <0,0001 |
|  |  | 594 | 2 | 0 | 0 | 0 | 0 |  |  |  |
|  | Globular | 32 | | 52 | | | | 84 |  |  |
|  |  | 18 | 14 | 0 | 4 | 46 | 2 |  |  |  |
|  | Dead | 70 | | 24 | | | | 94 |  |  |
|  |  | 30 | 40 | 0 | 0 | 14 | 10 |  |  |  |
|  | total | 698 | | 76 | | | | 774 |  |  |
|  |  | 642 | 56 | 0 | 4 | 60 | 12 |  |  |  |
| **14** | Vesicular | 636 | | 6 | | | | 642 | 522.9, 2 | <0,0001 |
|  |  | 632 | 4 | 2 | 0 | 4 | 0 |  |  |  |
|  | Globular | 102 | | 88 | | | | 190 |  |  |
|  |  | 64 | 38 | 20 | 0 | 62 | 6 |  |  |  |
|  | Dead | 22 | | 104 | | | | 126 |  |  |
|  |  | 14 | 8 | 0 | 0 | 36 | 68 |  |  |  |
|  | total | 760 | | 198 | | | | 958 |  |  |
|  |  | 710 | 50 | 22 | 0 | 102 | 74 |  |  |  |

Cell numbers in detailed and more general categories (Induced Pil1 / Non-induced Pil1) are indicated. Chi-squared tests for each timepoint were performed to determine the statistical significance of independence of the values. Chi-squared values and p-values for each time point are indicated. For the Chi-squared calculation the number of cells from the general categories was used.

**Table S3:** Probabilities of eisosome-expansion and/or assembly as predictive markers of Quiescence.

| **days PGD** | **mitochondrial morphology** | **Induced Pil1** | **Non induced Pil1** |
| --- | --- | --- | --- |
| **0** | Vesicular | 98,16849817 | 37,07865169 |
|  | Globular | 1,648351648 | 38,20224719 |
|  | Dead | 0,183150183 | 24,71910112 |
|  | total | 100 | 100 |
| **4** | Vesicular | 95,76659039 | 17,51824818 |
|  | Globular | 2,059496568 | 45,98540146 |
|  | Dead | 2,173913043 | 36,49635036 |
|  | total | 100 | 100 |
| **7** | Vesicular | 99,07100199 | 6,993006993 |
|  | Globular | 0,530856005 | 60,13986014 |
|  | Dead | 0,398142004 | 32,86713287 |
|  | total | 100 | 100 |
| **11** | Vesicular | 85,38681948 | 0 |
|  | Globular | 4,584527221 | 68,42105263 |
|  | Dead | 10,0286533 | 31,57894737 |
|  | total | 100 | 100 |
| **14** | Vesicular | 83,68421053 | 3,03030303 |
|  | Globular | 13,42105263 | 44,44444444 |
|  | Dead | 2,894736842 | 52,52525253 |
|  | total | 100 | 100 |
| **days PGD** | **mitochondrial morphology** | **Induced Pil1 and optimal assembly** | **Disassembly** |
| **0** | Vesicular | 98,71794872 | 64,07185629 |
|  | Globular | 1,282051282 | 22,15568862 |
|  | Dead | 0 | 13,77245509 |
|  | total | 100 | 100 |
| **4** | Vesicular | 95,73732719 | 20,97902098 |
|  | Globular | 2,073732719 | 44,05594406 |
|  | Dead | 2,188940092 | 34,96503497 |
|  | total | 100 | 100 |
| **7** | Vesicular | 99,06790945 | 10,13513514 |
|  | Globular | 0,532623169 | 58,10810811 |
|  | Dead | 0,399467377 | 31,75675676 |
|  | total | 100 | 100 |
| **11** | Vesicular | 92,52336449 | 1,515151515 |
|  | Globular | 2,803738318 | 50 |
|  | Dead | 4,672897196 | 48,48484848 |
|  | total | 100 | 100 |
| **14** | Vesicular | 89,01408451 | 4,032258065 |
|  | Globular | 9,014084507 | 50,80645161 |
|  | Dead | 1,971830986 | 45,16129032 |
|  | total | 100 | 100 |

**Probability % of cells with induced or non-induced Pil1, or with induced Pil1 and optimal eisosome assembly or eisosome disassembly, to be Quiescent, Senescent or Dead based on the morphology of their mitochondrial network.**

**Table S5: Statistical analysis of Figure 4F. Holm-Sidak's multiple comparisons test.**

| Time point | Comparisons | Summary | Adjusted P Value |
| --- | --- | --- | --- |
| G/2 | wt vs. mccΔ | ns | 0,8907 |
|  | wt vs. flpΔ | ns | 0,7586 |
|  | wt vs. coq3Δ | ns | 0,3648 |
|  | wt vs. atp1Δ | *** | 0,0004 |
|  | wt vs. wt + α-TOH | ns | 0,7726 |
|  | mccΔ vs. mccΔ + α-TOH | ns | 0,9951 |
|  | flpΔ vs. flpΔ + α-TOH | ns | 0,9981 |
|  | coq3Δ vs. coq3Δ + α-TOH | ns | 0,9984 |
|  | atp1Δ vs. atp1Δ + α-TOH | ns | 0,9984 |
|  | wt + α-TOH vs. mccΔ + α-TOH | ns | 0,2477 |
|  | wt + α-TOH vs. flpΔ + α-TOH | * | 0,0219 |
|  | wt + α-TOH vs. coq3Δ + α-TOH | * | 0,0127 |
|  | wt + α-TOH vs. atp1Δ + α-TOH | **** | <0,0001 |
| G/2 - GD | wt vs. mccΔ | ns | 0,9942 |
|  | wt vs. flpΔ | * | 0,0226 |
|  | wt vs. coq3Δ | ** | 0,0013 |
|  | wt vs. atp1Δ | **** | <0,0001 |
|  | wt vs. wt + α-TOH | ** | 0,0043 |
|  | mccΔ vs. mccΔ + α-TOH | ns | 0,8926 |
|  | mccΔ vs. flpΔ + α-TOH | ns | 0,9942 |
|  | flpΔ vs. flpΔ + α-TOH | ** | 0,0049 |
|  | coq3Δ vs. coq3Δ + α-TOH | ns | 0,6609 |
|  | atp1Δ vs. atp1Δ + α-TOH | ns | 0,9942 |
|  | wt + α-TOH vs. mccΔ + α-TOH | ns | 0,1408 |
|  | wt + α-TOH vs. flpΔ + α-TOH | * | 0,0199 |
|  | wt + α-TOH vs. coq3Δ + α-TOH | **** | <0,0001 |
|  | wt + α-TOH vs. atp1Δ + α-TOH | **** | <0,0001 |
| GD | wt vs. mccΔ | ns | 0,7307 |
|  | wt vs. flpΔ | * | 0,0222 |
|  | wt vs. coq3Δ | *** | 0,0001 |
|  | wt vs. atp1Δ | **** | <0,0001 |
|  | wt vs. wt + α-TOH | ns | 0,8211 |
|  | mccΔ vs. mccΔ + α-TOH | ns | 0,7307 |
|  | flpΔ vs. flpΔ + α-TOH | ns | 0,9659 |
|  | coq3Δ vs. coq3Δ + α-TOH | ns | 0,8563 |
|  | atp1Δ vs. atp1Δ + α-TOH | ns | 0,8802 |
|  | wt + α-TOH vs. mccΔ + α-TOH | ns | 0,8211 |
|  | wt + α-TOH vs. flpΔ + α-TOH | ** | 0,0023 |
|  | wt + α-TOH vs. coq3Δ + α-TOH | *** | 0,0009 |
|  | wt + α-TOH vs. atp1Δ + α-TOH | **** | <0,0001 |
| PGD 1 day | wt vs. mccΔ | ns | 0,6294 |
|  | wt vs. flpΔ | *** | 0,0006 |
|  | wt vs. coq3Δ | **** | <0,0001 |
|  | wt vs. atp1Δ | **** | <0,0001 |
|  | wt vs. wt + α-TOH | ns | 0,8121 |
|  | mccΔ vs. mccΔ + α-TOH | ns | 0,6453 |
|  | flpΔ vs. flpΔ + α-TOH | ns | 0,122 |
|  | coq3Δ vs. coq3Δ + α-TOH | ns | 0,9481 |
|  | atp1Δ vs. atp1Δ + α-TOH | ns | 0,8121 |
|  | wt + α-TOH vs. mccΔ + α-TOH | ns | 0,8121 |
|  | wt + α-TOH vs. flpΔ + α-TOH | ns | 0,0761 |
|  | wt + α-TOH vs. coq3Δ + α-TOH | **** | <0,0001 |
|  | wt + α-TOH vs. atp1Δ + α-TOH | **** | <0,0001 |
| PGD 1.5 days | wt vs. mccΔ | ns | 0,4149 |
|  | wt vs. flpΔ | ** | 0,0013 |
|  | wt vs. coq3Δ | **** | <0,0001 |
|  | wt vs. atp1Δ | **** | <0,0001 |
|  | wt vs. wt + α-TOH | ns | 0,1897 |
|  | mccΔ vs. mccΔ + α-TOH | ns | 0,3349 |
|  | flpΔ vs. flpΔ + α-TOH | *** | 0,0005 |
|  | coq3Δ vs. coq3Δ + α-TOH | ns | 0,8481 |
|  | atp1Δ vs. atp1Δ + α-TOH | ns | 0,9984 |
|  | wt + α-TOH vs. mccΔ + α-TOH | ns | 0,2795 |
|  | wt + α-TOH vs. flpΔ + α-TOH | ns | 0,3025 |
|  | wt + α-TOH vs. coq3Δ + α-TOH | **** | <0,0001 |
|  | wt + α-TOH vs. atp1Δ + α-TOH | **** | <0,0001 |
| PGD 2 days | wt vs. mccΔ | * | 0,0396 |
|  | wt vs. flpΔ | **** | <0,0001 |
|  | wt vs. coq3Δ | **** | <0,0001 |
|  | wt vs. atp1Δ | **** | <0,0001 |
|  | wt vs. wt + α-TOH | ns | 0,3827 |
|  | mccΔ vs. mccΔ + α-TOH | ns | 0,0894 |
|  | flpΔ vs. flpΔ + α-TOH | *** | 0,0008 |
|  | coq3Δ vs. coq3Δ + α-TOH | ns | 0,6479 |
|  | atp1Δ vs. atp1Δ + α-TOH | ns | 0,9356 |
|  | wt + α-TOH vs. mccΔ + α-TOH | ns | 0,2747 |
|  | wt + α-TOH vs. flpΔ + α-TOH | ns | 0,0664 |
|  | wt + α-TOH vs. coq3Δ + α-TOH | **** | <0,0001 |
|  | wt + α-TOH vs. atp1Δ + α-TOH | **** | <0,0001 |
| PGD 3 days | wt vs. mccΔ | * | 0,0377 |
|  | wt vs. flpΔ | **** | <0,0001 |
|  | wt vs. coq3Δ | **** | <0,0001 |
|  | wt vs. atp1Δ | **** | <0,0001 |
|  | wt vs. wt + α-TOH | ns | 0,2787 |
|  | mccΔ vs. mccΔ + α-TOH | ns | 0,1072 |
|  | flpΔ vs. flpΔ + α-TOH | *** | 0,0007 |
|  | coq3Δ vs. coq3Δ + α-TOH | ns | 0,5298 |
|  | atp1Δ vs. atp1Δ + α-TOH | ns | 0,8395 |
|  | wt + α-TOH vs. mccΔ + α-TOH | ns | 0,1292 |
|  | wt + α-TOH vs. flpΔ + α-TOH | ns | 0,1072 |
|  | wt + α-TOH vs. coq3Δ + α-TOH | **** | <0,0001 |
|  | wt + α-TOH vs. atp1Δ + α-TOH | **** | <0,0001 |
| PGD 4 days | wt vs. mccΔ | **** | <0,0001 |
|  | wt vs. flpΔ | **** | <0,0001 |
|  | wt vs. coq3Δ | **** | <0,0001 |
|  | wt vs. atp1Δ | **** | <0,0001 |
|  | wt vs. wt + α-TOH | ns | 0,7633 |
|  | mccΔ vs. mccΔ + α-TOH | * | 0,0112 |
|  | flpΔ vs. flpΔ + α-TOH | **** | <0,0001 |
|  | coq3Δ vs. coq3Δ + α-TOH | ns | 0,3978 |
|  | atp1Δ vs. atp1Δ + α-TOH | ns | 0,8476 |
|  | wt + α-TOH vs. mccΔ + α-TOH | ns | 0,171 |
|  | wt + α-TOH vs. flpΔ + α-TOH | ns | 0,091 |
|  | wt + α-TOH vs. coq3Δ + α-TOH | **** | <0,0001 |
|  | wt + α-TOH vs. atp1Δ + α-TOH | **** | <0,0001 |
| PGD 5 days | wt vs. mccΔ | ns | 0,3997 |
|  | wt vs. flpΔ | **** | <0,0001 |
|  | wt vs. coq3Δ | **** | <0,0001 |
|  | wt vs. atp1Δ | **** | <0,0001 |
|  | wt vs. wt + α-TOH | ** | 0,0029 |
|  | mccΔ vs. mccΔ + α-TOH | ns | 0,1163 |
|  | flpΔ vs. flpΔ + α-TOH | **** | <0,0001 |
|  | coq3Δ vs. coq3Δ + α-TOH | ns | 0,3556 |
|  | atp1Δ vs. atp1Δ + α-TOH | ns | 0,8287 |
|  | wt + α-TOH vs. mccΔ + α-TOH | * | 0,0264 |
|  | wt + α-TOH vs. flpΔ + α-TOH | ** | 0,009 |
|  | wt + α-TOH vs. coq3Δ + α-TOH | **** | <0,0001 |
|  | wt + α-TOH vs. atp1Δ + α-TOH | **** | <0,0001 |
| PGD 6 days | wt vs. mccΔ | **** | <0,0001 |
|  | wt vs. flpΔ | **** | <0,0001 |
|  | wt vs. coq3Δ | **** | <0,0001 |
|  | wt vs. atp1Δ | **** | <0,0001 |
|  | wt vs. wt + α-TOH | ns | 0,4027 |
|  | mccΔ vs. mccΔ + α-TOH | * | 0,01 |
|  | flpΔ vs. flpΔ + α-TOH | **** | <0,0001 |
|  | coq3Δ vs. coq3Δ + α-TOH | ** | 0,0026 |
|  | atp1Δ vs. atp1Δ + α-TOH | ns | 0,9124 |
|  | wt + α-TOH vs. mccΔ + α-TOH | * | 0,0367 |
|  | wt + α-TOH vs. flpΔ + α-TOH | ns | 0,0878 |
|  | wt + α-TOH vs. coq3Δ + α-TOH | **** | <0,0001 |
|  | wt + α-TOH vs. atp1Δ + α-TOH | **** | <0,0001 |

**Table S6: List of transitions used for targeted lipidomics.**

| **Description** | **Q1 mass** | **Q2 mass** | **CE (V)** | **CXP V** | **DP V** | **EP V** |
| --- | --- | --- | --- | --- | --- | --- |
| Ceramide d17:1/24:0 +HCOO | 680,610 | 392,400 | -56 | 15 | -35 | -10 |
| 16:0_18:1 | 716,523 | 255,232 | -80 | 5 | -120 | -10 |
| 16:1_18:1 | 714,523 | 281,249 | -80 | 5 | -120 | -10 |
| 16:1_18:2 | 712,523 | 279,249 | -80 | 5 | -120 | -10 |
| 16:1_18:3 | 710,523 | 277,249 | -80 | 5 | -120 | -10 |
| 18:1_18:1 | 742,55 | 281,249 | -80 | 5 | -120 | -10 |
| 18:1_18:2 | 740,55 | 279,249 | -80 | 5 | -120 | -10 |
| 18:1_18:3 | 738,55 | 277,249 | -80 | 5 | -120 | -10 |
| 18:2_18:2 | 738,55 | 279,249 | -80 | 5 | -120 | -10 |
| 18:3_18:3 | 734,55 | 277,249 | -80 | 5 | -120 | -10 |
| 16:1_18:1<O> | 730,523 | 297,249 | -80 | 5 | -120 | -10 |
| 16:1_18:2<O> | 728,523 | 295,249 | -80 | 5 | -120 | -10 |
| 16:1_18:3<O> | 726,523 | 293,249 | -80 | 5 | -120 | -10 |
| 16:1_18:1<2O> | 746,523 | 313,249 | -80 | 5 | -120 | -10 |
| 16:1_18:2<2O> | 744,523 | 311,249 | -80 | 5 | -120 | -10 |
| 16:1_18:3<2O> | 742,523 | 309,249 | -80 | 5 | -120 | -10 |
| 18:1_18:1<O> | 758,55 | 281,249 | -80 | 5 | -120 | -10 |
| 18:1_18:2<O> | 756,55 | 279,249 | -80 | 5 | -120 | -10 |
| 18:1_18:3<O> | 754,55 | 277,249 | -80 | 5 | -120 | -10 |
| 18:2_18:2<O> | 754,55 | 295,249 | -80 | 5 | -120 | -10 |
| 18:3_18:3<O> | 750,55 | 277,249 | -80 | 5 | -120 | -10 |
| 18:1_18:1<2O> | 774,55 | 281,249 | -80 | 5 | -120 | -10 |
| 18:2_18:1<2O> | 772,55 | 279,249 | -80 | 5 | -120 | -10 |
| 18:3_18:1<2O> | 770,55 | 277,249 | -80 | 5 | -120 | -10 |
| 18:1_18:2<2O> | 772,55 | 311,249 | -80 | 5 | -120 | -10 |
| 18:1_18:3<2O> | 770,55 | 309,249 | -80 | 5 | -120 | -10 |
| 18:2_18:2<2O> | 770,55 | 279,249 | -80 | 5 | -120 | -10 |
| 18:3_18:3<2O> | 766,55 | 277,249 | -80 | 5 | -120 | -10 |

**Table S7. *S. cerevisiae* strains and plasmids used in this study.**

| **Strain** | ***Genotype*** | **Reference or source** |
| --- | --- | --- |
| Sigma1278b |  | Laboratory collection |
| 23344c | *ura3* | Laboratory collection |
| AM029 | *ura3::PIL1-4A-GFP~~-~~URA3 LSP1-mCHERRY-KANMX6 nce102Δ::HPHMX4 pil1Δ ura3* | This study |
| AM042 | *PIL1-GFP-CaURA3 LSP1-mCHERRY-KANMX6 ura3* | This study |
| AM043 | *LSP1-mCHERRY-KANMX6 pil1Δ ura3* | This study |
| AM045 | *ura3::PIL1-4A-GFP~~-~~URA3 LSP1-mCHERRY-KANMX6 pil1Δ ura3* | This study |
| AM101 | *fmp45Δ::HPHMX4 PIL1-mCHERRY-KANMX6 URA3* | This study |
| AM105 | *pst2Δ::HPHMX4 PIL1-mCHERRY-KANMX6 URA3* | This study |
| AM115 | *pun1Δ::KiURA3 PIL1-mCHERRY-KANMX6 ura3* | This study |
| AM117 | *pil1Δ URA3* | This study |
| AM119 | *nce102Δ::KiURA3 PIL1-mCHERRY-KANMX6 ura3* | This study |
| AM121 | *PIL1-mCHERRY-KANMX6 URA3* | This study |
| AM129 | *Lsp1-GFP-CaURA3 pil1Δ ura3* | This study |
| AM139 | *LSP1-GFP-CaURA3 PIL1-mCHERRY-KANMX6 ura3* | This study |
| AM145 | *ILV3-GFP-CaURA3 PIL1-mCHERRY-KANMX6 ura3* | This study |
| AM164, AM165 | *lsp1Δ::HPHMX4 pil1Δ ura3* | This study |
| AM170 | *lsp1Δ::HPHMX4 PST2-GFP-CaURA3 pil1Δ ura3* | This study |
| AM174 | *YNL194C-mCHERRY-KANMX6 LSP1-GFP-URA3 pil1Δ ura3* | This study |
| AM186 | *lsp1Δ::HPHMX4 pil1Δ URA3* | This study |
| AM203 | *lsp1Δ::HPHMX4 ILV3-GFP-CaURA3 pil1Δ ura3* | This study |
| AM220 | *lsp1Δ::HPHMX4* | This study |
| AM237 | *nce102Δ::HPHMX4 PIL1-GFP-CaURA3 LSP1-mCHERRY-KANMX6 ura3* | This study |
| AM238 | *nce102Δ::HPHMX4* | This study |
| AM254 | *atp1Δ::HPHMX4 ILV3-GFP-CaURA3 PIL1-mCHERRY-KANMX6 ura3* | This study |
| AM282 | *rho0 ILV3-GFP-CaURA3 PIL1-mCHERRY-KANMX6 ura3* | This study |
| AM288 | *cox7Δ::HPHMX4 ILV3-GFP-CaURA3 PIL1-mCHERRY-KANMX6 ura3* | This study |
| AM290 | *cbs1Δ::HPHMX4 ILV3-GFP-CaURA3 PIL1-mCHERRY-KANMX6 ura3* | This study |
| AM292 | *atp1Δ::HPHMX4 ura3* | This study |
| AM294 | *atp1Δ::HPHMX4* | This study |
| AM296 | *PST2-GFP-CaURA3 PIL1-mCHERRY-KANMX6 ura3* | This study |
| AM317 | *gpx1Δ::KANMX6 gpx3Δ::KiURA3 gpx2Δ rfs1Δ ycp4Δ::NATMX4 pst2Δ::HPHMX4 ura3* | This study |
| AM323 | *gpx3Δ::KiURA3 gpx1Δ gpx2Δ rfs1Δ ycp4Δ::NATMX4 pst2Δ::HPHMX4 ura3* | This study |
| AM325 | *cqd1Δ::KANMX6 gpx3Δ::KiURA3 gpx1Δ gpx2Δ rfs1Δ ycp4Δ::NATMX4 pst2Δ::HPHMX4 ura3* | This study |
| AM345 | *cqd1Δ::HPHMX4 gpx2Δ::KiURA3 gpx1Δ::KANMX6 gpx3Δ::NATMX4 ura3* | This study |
| AM349 | *LSP1-GFP-KANMX6 nce102Δ::HPHMX4* | This study |
| CG116 | *lsp1Δ::HPHMX4 PIL1-mCHERRY-KANMX6 can1Δ gap1Δ ura3* | Gournas et al., 2018 |
| CG210 | *ynl194cΔ::KiURA3 PIL1-mCHERRY-KANMX6 ura3* | This study |
| CG254 | *PAB1-GFP-CaURA3 PIL1-mCHERRY-KANMX6 ura3* | This study |
| CG271 | *rfs1Δ::KANMX6 ycp4Δ::NATMX4 pst2Δ::KANMX6 URA3* | This study |
| CG272 | *RFS1-GFP-CaURA3 PIL1-mCHERRY-KANMX6 ura3* | This study |
| CG277 | *coq3Δ::HPHMX4* | This study |
| CG280 | *rfs1Δ::KANMX6 ycp4Δ::NATMX4 pst2Δ::HPHMX4 ura3* | This study |
| CG281 | *YCP4-GFP-KiURA3 PIL1-mCHERRY-KANMX6 ura3* | This study |
| CG282 | *RFS1-GFP-KANMX6 lsp1Δ::HPHMX4 pil1Δ URA3* | This study |
| CG309 | *YCP4-GFP-CaURA3 lsp1Δ::HPHMX4 pil1Δ ura3* | This study |
| CG317 | *FTR1-GFP-CaURA3 PIL1-mCHERRY-KANMX6 ura3* | This study |
| CG319 | *FTR1-GFP-CaURA3 PIL1-mCHERRY-KANMX6 rfs1Δ ycp4Δ::NATMX4 pst2Δ::HPHMX4 ura3* | This study |
| CG371 | *gpx1Δ::HPHMX4 gpx3Δ::NATMX4 gpx2Δ:: KANMX6* | This study |
| GK022 | *PIL1-mCHERRY-KANMX6 ura3* | This study |
| SG035 | *PIL1-mCHERRY-KANMX6 can1Δ gap1Δ ura3* | Gournas et al., 2018 |
| **Plasmid** | **Description** | **Reference or source** |
| pCG118 | CEN-ARS NCE102-5GA-GFP-URA3 | Gournas et al., 2018 |
| pCG140 | CEN-ARS CAN1-(K5R/K13R/K42R/K45R/K47R/K85R/K89R)-GFP-URA3 | This study |
| pFL038 | CEN-ARS-URA3 | Gournas et al., 2018 |
| pAM3 | CEN-ARS-pMET25-PRESU9-GFP-pADH1-PRECOX4-mCHERRY-URA3 | This study |
| pAM1 | CEN-ARS PIL1-URA3 | This study |
| pCG142 | CEN-ARS LSP1-5GA-GFP-URA3 | This study |
| pGFP-Atg8 URA3 | CEN-ARS GFP-ATG8-URA3 | Laboratory collection |
