## Supplementary material for "Ferroptosis-Protective Membrane Domains in Quiescence": Table S8

### General Lipidomics Workflow

#### Overall study design

| | | | |
|---|---|---|---|
| Title of the study | Determination of peroxidised lipid species in Quiescent cells of Saccharomyces cerevisiae | | |
| Principle investigator | Florian Fröhlich | | |
| Institution | Department of Biology/Chemistry, Molecular Membrane Biology Group, University of Osnabrück | | |
| Corresponding Email | | | |
| Document creation date | 11/14/2022 | Clinical | No |
| Is the workflow targeted or untargeted? | Targeted | | |

#### Lipid extraction

| | | | |
|---|---|---|---|
| Extraction method | 2-phase system | 2-phase system | 2-step extraction |
| pH adjustment | None | Were internal standards added prior extraction? | Yes |

#### Analytical platform

| | | | |
|---|---|---|---|
| MS type | QTrap | MS Level | MS2 |
| MS vendor | SCIEX | Mass window for precursor ion isolation (in Da total isolation window) | 1 |
| Ion source | ESI | Mass resolution for detected ion at MS2 | Low resolution |
| Direct type | Syringe | Resolution in Da at MS2 | 1 |

#### Quality control

| | | | |
|---|---|---|---|
| Blanks | No | Quality control | No |

#### Method qualification and validation

| | | | |
|---|---|---|---|
| Method validation | Yes | Precison | No |
| Lipid recovery | No | Accuracy | No |
| Dynamic quantification range | Yes | Guidelines followed | None |
| Limit of quantitation (LOQ)/Limit of detection (LOD) | No | | |

#### Reporting

| | | | |
|---|---|---|---|
| Are reported raw data uploaded into repository? | Available on request | Raw data upload | Available on request |
